# Human iPSCs and iPSC-Derived Endothelial Cells from PAD Patients and Healthy Donors Exhibit Comparable Characteristics and Potency: Implications for Autologous Cell Therapy in Peripheral Artery Disease

**DOI:** 10.64898/2026.09.04.749119

**Authors:** Jung Yoon Bae, Shin-Jeong Lee, Seongho Bae, Juhyun Park, Jihyeoun Jung, Ji Woong Han, Cholomi Jung, Sinhye Park, Yonghak Kim, Jungwoo Kim, Hyun-Kyung Kim, Seung-Jun Lee, Young-Guk Ko, Donghoon Choi, Hyun Ok Kim, Young-sup Yoon

**Author notes:** Correspondence to: Young-sup Yoon, MD, PhD Division of Cardiology, Department of Medicine, Emory University School of Medicine, 275 Haygood Drive, HSRBII N248, Atlanta, GA, 30322.

## Abstract

**BACKGROUND:** Peripheral artery disease (PAD) can lead to amputation in advanced cases, making cell therapy using human induced pluripotent stem cells (hiPSCs) a promising therapeutic option. hiPSC-derived endothelial cells (hiPSC-ECs) have shown favorable effects in treating experimental ischemic cardiovascular disease. An autologous approach for PAD patients is preferable to avoid immunological reactions. However, it is yet unknown whether hiPSCs and hiPSC-ECs derived from PAD patients have similar characteristics and potency compared to those derived from non-PAD donors. Therefore, we explored whether there are significant differences in the characteristics and potency of hiPSCs and hiPSC-ECs between non-PAD donors and PAD patients.

**METHODS:** We generated hiPSCs from the blood of non-PAD donors and patients with PAD. We determined the pluripotency of hiPSCs using qRT-PCR, flow cytometry, immunostaining, RNA-seq, and teratoma formation assay. The genetic stability of hiPSCs was confirmed by G-banding karyotyping. We then differentiated hiPSCs into endothelial cells (ECs) in a clinically compatible manner. We compared the characteristics and therapeutic potency of hiPSC-ECs derived from non-PAD and PAD groups using qRT-PCR, flow cytometry, immunostaining, and RNA-seq. In vitro endothelial functions were determined by nitric oxide (NO) production and capillary network formation. Therapeutic potency and incorporation of engrafted hiPSC-ECs were evaluated in a hindlimb ischemia model.

**RESULTS:** We successfully generated hiPSCs from the blood of seven non-PAD donors and eight PAD patients. Both non-PAD and PAD-derived hiPSCs exhibited similar expression levels of pluripotency markers. All hiPSCs, regardless of the group, formed teratomas and showed normal karyotypes. RNA-seq analyses revealed similar gene expression profiles between the groups. hiPSC-ECs derived from both non-PAD and PAD donors exhibited similar expression levels of EC markers at both the gene and protein levels. RNA-seq analyses showed no significant overall differences in gene expression profiles between the groups. Functional analyses demonstrated similar endothelial characteristics and function in hiPSC-ECs from both groups. Both groups showed similar perfusion recovery, limb salvage, and vessel-forming capacity. Engrafted hiPSC-ECs from both groups also exhibited similar angiogenic and vessel-forming capabilities.

**CONCLUSIONS:** Our study demonstrated no significant differences in hiPSCs and hiPSC-ECs derived from non-PAD donors and PAD patients in terms of molecular and cell biological characteristics, therapeutic effects, and vessel-forming capability. Our study indicates that hiPSCs and hiPSC-ECs derived from PAD patients can serve as a novel platform for autologous cell therapy.

## INTRODUCTION

Peripheral artery disease (PAD) is characterized by atherosclerotic plaque accumulation along the walls of peripheral arteries in the arms and legs. A significant proportion of patients progress to chronic limb-threatening ischemia (CLTI), which manifests as rest pain, non-healing ulcers, and gangrene. Approximately 40% of CLTI patients who fail conventional treatment such as medication, endovascular intervention, or surgery undergo major amputation within one year, with a substantial decline in quality of life.^1–3^ Patients with advanced PAD or CLTI therefore require innovative approaches, such as cell-based therapy, to prevent limb loss.

Earlier studies employed adult stem or progenitor cells, including endothelial progenitor cells, mesenchymal stem cells, and bone marrow mononuclear cells, to induce neovascularization.^4, 5^ Although preclinical studies were promising, clinical trial outcomes have been modest at best,^6–8^ largely because these cells fail to form durable vessels in vivo and rely primarily on paracrine effects.^9–11^ Human pluripotent stem cells (hPSCs), which include human embryonic stem cells (hESCs) and human induced pluripotent stem cells (hiPSCs) have therefore emerged as a more compelling option for vascular regeneration, given their pluripotency, proliferative capacity, and robust potential to differentiate into target lineages.^12–15^

While hESCs have shown experimental promise, their clinical application is constrained by ethical and immunogenic concerns. hiPSCs exhibit cellular and molecular characteristics comparable to hESCs but avoid these constraints and uniquely enable autologous therapy.^15^ Accordingly, numerous studies have differentiated hiPSCs into therapeutically relevant lineages.^16–20^ For ischemic cardiovascular diseases (CVD), hiPSC-derived endothelial cells (hiPSC-ECs) have been shown to contribute to vessel formation in vivo and to ameliorate animal models through both angiogenesis and vasculogenesis.^21–26^ Several groups have generated hiPSC-ECs under clinically compatible conditions, highlighting their translational potential.^27, 28^ We recently demonstrated that hiPSC-ECs differentiated under fully defined conditions produce sustained vessel-forming effects in ischemic hindlimbs for more than 10 months,^26^ supporting their promise for treating ischemic CVD.

Autologous therapy is generally preferred over allogeneic approaches because it avoids immunological reactions. However, PAD patients frequently carry comorbid conditions such as diabetes, hypertension, and hypercholesterolemia. Before patient-derived hiPSC-ECs can be advanced to clinical application, it is therefore essential to determine whether they retain the characteristics and therapeutic potency of hiPSC-ECs derived from non-PAD donors. To date, most studies of patient-specific hiPSCs and their derivatives have focused on disease modeling for monogenic or inherited diseases.^29–38^ hiPSCs from such patients retain the causal genotype,^37–39^ and pathogenic mutations are preserved upon lineage-directed differentiation.^29, 34, 40^ In contrast, hiPSCs from patients with non-genetic diseases generally display characteristics comparable to those of healthy controls,^41, 42^ suggesting that disease signatures are largely erased during reprogramming. Findings on differentiated derivatives, however, have been mixed: some studies report no differences between patient and control groups,^42–45^ whereas others describe altered phenotypes or functional deficits.^32, 46^ Importantly, these studies were limited by small sample sizes or by reliance on in vitro assays alone, restricting interpretation of their findings.

Here, we addressed this gap through a large-scale, integrated comparison of hiPSCs and hiPSC-ECs derived from eight PAD patients and seven non-PAD donors at the molecular, cellular, and in vivo functional levels, using a clinically compatible differentiation protocol. We found that hiPSCs and hiPSC-ECs from PAD patients were molecularly, cellularly, and functionally equivalent to those from non-PAD donors, and that hiPSC-ECs from both groups exhibited comparable therapeutic potency in a murine hindlimb ischemia model. To our knowledge, this is the first such head-to-head comparison. Together, these findings provide a foundation for autologous cell therapy using patient-derived hiPSC-ECs in PAD.

## METHODS

### Human Subjects

All human studies were conducted in accordance with the guidelines and approval of the Institutional Review Board (IRB) of Yonsei University Severance Hospital, Seoul, Korea (IRB No. 4-2016-0815). Written informed consent was obtained from all participants prior to enrollment. Peripheral blood samples were collected from patients with peripheral artery disease (PAD) and from non-PAD donors frequency-matched to the PAD patients by age at Yonsei University Severance Hospital.

### Animal Studies

All animal experiments were performed in accordance with the *Guidelines for the Care and Use of Laboratory Animals* and were approved by the Institutional Animal Care and Use Committees (IACUCs) of Emory University and Yonsei University College of Medicine (IACUC No. 2017-0165). Male and female immunodeficient BALB/c nude mice (6–8 weeks old) were maintained under specific pathogen–free (SPF) conditions at both institutions.

### hiPSC Generation

Human induced pluripotent stem cells (hiPSCs) were generated from peripheral blood mononuclear cells (PBMCs) obtained from non-PAD and PAD donors using episomal vectors (Epi5™ Episomal iPSC Reprogramming Kit, A15960, Invitrogen, Waltham, USA).^47^ Peripheral blood was collected from seven non-PAD donors over 50 years of age and eight PAD patients classified as Rutherford categories 4-5. PBMCs were isolated using Lymphoprep (StemCell Technologies, Oslo, Norway) and cultured in erythroid expansion medium to induce erythroid precursor proliferation. Electroporation was performed on erythroblasts with three episomal plasmids encoding the five reprogramming factors-*OCT4, SOX2, KLF4, L-MYC,* and *LIN28*-using a NEPA21 electroporator (Nepa Gene Co., Ltd., Ichikawa, Japan). Transfected cells were seeded on Matrigel-coated plates and maintained in erythroid expansion medium on day 2, followed by ReproTeSR Basal Medium (StemCell Technologies) on days 3 and 5 post-transfection. From day 7 onward, the medium was replaced daily with complete ReproTeSR medium. By approximately day 20, emerging colonies exhibited morphology typical of human embryonic stem cells (hESCs). For clonal expansion, colonies were manually picked and transferred using a feeder-free system. Cells were cultured on Vitronectin XF–coated plates (StemCell Technologies) in mTeSR™1 medium (StemCell Technologies) at 37°C in a humidified incubator with 5% CO₂. hiPSCs were maintained in mTeSR1 medium with daily medium changes until reaching 80–90% confluence. Newly established hiPSCs (up to passage 5) were manually passaged using drawn glass Pasteur pipettes, while enzymatic passaging with ReLeSR (StemCell Technologies) was performed from passage 5 onward. Cells were subcultured every 5–7 days.

### Differentiation of hiPSCs into Endothelial Cells (ECs)

Directed differentiation of hiPSCs into endothelial cells (ECs) was performed as previously described.^26^ Dissociated hiPSC clumps (passages 15–25) were seeded onto 0.01% collagen-coated plates and cultured in DMEM/F12 (1:1) supplemented with 20% KnockOut™ Serum Replacement (Thermo Fisher Scientific) with or without the addition of lineage-specific differentiation factors. Cells were maintained under these conditions for 14–16 days to induce endothelial lineage commitment. The resulting differentiating cells were then dissociated using 0.25% Trypsin-EDTA (Gibco) and replated onto collagen-coated plates for further expansion and were passaged. All experiments were performed using hiPSC-ECs at passage 2 or later.

### Quantitative Real-Time PCR (qRT–PCR)

Total RNA was extracted from hiPSCs or hiPSC-derived endothelial cells (hiPSC-ECs) using TRIsure™ reagent (BIO-38032, Bioline Ltd., London, UK). Complementary DNA (cDNA) was synthesized from purified RNA using the SensiFast™ cDNA Synthesis Kit (BIO-65053, Bioline Ltd.) according to the manufacturer’s instructions. Quantitative real-time PCR was performed using gene-specific primers and probes (see Table S2) on an ABI PRISM 7500 Sequence Detection System (Applied Biosystems, Waltham, USA). Relative mRNA expression levels were normalized to *GAPDH* expression.

### Flow Cytometry

Flow cytometry was performed as previously described.^48, 49^ Cells were resuspended in MACS Separation Buffer (130-091-221, Miltenyi Biotec, Bergisch Gladbach, Germany) containing 0.05% bovine serum albumin (BSA; 130-091-376, Miltenyi Biotec) and incubated for 20 min at 4°C with directly conjugated PE-, APC-, or FITC-labeled antibodies. Corresponding isotype-matched IgGs served as negative controls. The following antibodies were used: TRA-1-60 (560193, BD Biosciences, Franklin Lakes, USA), SSEA4 (MC-813-70, BioLegend, San Diego, USA), CDH5/VE-cadherin/CD144 (17-1449-42, eBioscience™, San Diego, USA), KDR (FAB357P, R&D Systems, Minneapolis, USA), PECAM1/CD31 (130-098-173, Miltenyi Biotec), TEK (FAB3131P, R&D Systems), and VWF (ab8822, Abcam, Cambridge, UK). Flow-cytometric data were acquired using an Accuri C6 Flow Cytometer (BD Biosciences) and analyzed with FlowLogic software (Miltenyi Biotec) using appropriate isotype controls.

### Immunostaining

Immunostaining was performed as previously described.^48, 49^ Cells were washed with phosphate-buffered saline (PBS), fixed in 4% paraformaldehyde (PFA), and incubated with the following primary antibodies: OCT4 (1:100, A24867, Thermo Fisher Scientific, Waltham, USA), SSEA4 (1:100, A24866, Thermo Fisher Scientific), SOX2 (1:100, A24759, Thermo Fisher Scientific), TRA-1-60 (1:100, A24868, Thermo Fisher Scientific), CDH5 (1:200, sc-28644, Santa Cruz Biotechnology, Dallas, USA), and PECAM1 (1:100, CBL-468, Millipore, Burlington, USA). After primary antibody incubation, cells were treated with Alexa Fluor 488– or Alexa Fluor 594–conjugated secondary antibodies (1:250, Thermo Fisher Scientific; 1:1000, BD Biosciences). Nuclei were counterstained with DAPI where indicated. Immunofluorescence images were acquired using a Nikon fluorescence microscope or a Zeiss LSM 700/780 confocal laser scanning microscope (Carl Zeiss).

### Teratoma Formation Assay

A total of 1 × 10⁷ hiPSCs were injected subcutaneously into both flanks of 7- to 9-week-old BALB/c nude mice (Central Lab. Animal Inc., Seoul, Korea). Teratomas developed within 6–12 weeks and were harvested for histological analysis. Excised tissues were fixed, embedded in paraffin, sectioned, and stained with hematoxylin and eosin (H&E). Images were acquired using a BX53 upright microscope (Olympus, Tokyo, Japan).

### Analysis of Chromosomal Abnormalities

Chromosomal integrity of hiPSC lines was assessed using standard G-banding karyotype analysis.^50^ Cells were fixed and processed according to established protocols, and analyses were performed by GenDix (Seoul, Korea). Twenty-three pairs of human chromosomes were examined for each hiPSC line.

### Bulk RNA Sequencing

Total RNA was extracted from two biological replicates for each donor-derived line using the miRNeasy Mini Kit (QIAGEN) according to the manufacturer’s protocol. RNA integrity and concentration were assessed using an Agilent 2100 BioAnalyzer (Agilent Technologies, Santa Clara, USA), and only samples with an RNA Integrity Number (RIN) greater than 8 were used for sequencing. Polyadenylated [poly(A)] mRNA was enriched using magnetic oligo(dT) beads and subsequently fragmented. First- and second-strand cDNA synthesis, end repair, A-tailing, and adapter ligation were performed using the TruSeq Stranded mRNA Sample Preparation Kit (Illumina, San Diego, USA). Libraries with insert sizes ranging from 120–200 base pairs (bp) were sequenced on an Illumina NovaSeq 6000 platform, generating 150-bp paired-end reads with an average depth of ∼27 million reads per library. Raw reads were processed for quality control, and only high-quality (clean) reads were used for downstream analysis.

### Bioinformatics Analysis

Raw sequencing reads were pre-processed using Cutadapt to remove adapter sequences and low-quality bases.^51^ Filtered reads were aligned to the human reference genome (Genome Reference Consortium Human Build 38; GRCh38/hg38)^52^ using STAR.^53^ Gene-level expression quantification was performed using featureCounts with default parameters.^54^ Differential expression analysis between groups was conducted using DESeq2.^55^ Multiple testing correction was performed using the Benjamini– Hochberg false discovery rate (FDR) method. Differentially expressed genes (DEGs) were identified using a Benjamini–Hochberg-adjusted p value < 0.05. Normalized gene expression values generated by DESeq2 were used for downstream unsupervised analyses, including principal component analysis (PCA), hierarchical clustering, Pearson correlation analysis, and K-means clustering. Additional transcriptomic analyses were performed using iDEP v0.91 (http://bioinformatics.sdstate.edu/idep/).^56^ Pearson correlation coefficients were calculated using normalized expression values. For K-means clustering, the top differentially expressed genes between hiPSCs and hiPSC-ECs were grouped according to their expression patterns across samples. Gene Ontology (GO) Biological Process enrichment analysis was subsequently performed for each gene cluster using gene sets from the Gene Ontology database implemented in iDEP. For gene set enrichment analysis (GSEA), genes ranked according to differential expression between hiPSCs and hiPSC-ECs were analyzed using Gene Set Enrichment Analysis software (v4.1.0; Broad Institute) with 1,000 gene set permutations, no dataset collapse, and weighted enrichment statistics.^57^ Gene sets with a false discovery rate (FDR) q value < 0.25 and nominal p value < 0.05 were considered significantly enriched.

### Intracellular Nitric Oxide Detection

Intracellular nitric oxide (NO) production was assessed using 4-amino-5-methylamino-2′,7′-difluorofluorescein diacetate (DAF-FM diacetate; Invitrogen) as previously described.^58^ hiPSC-ECs were incubated with DAF-FM diacetate for 1 h at 37°C. After washing to remove excess probe, cells were incubated for an additional 20 min to allow complete de-esterification of intracellular DAF-FM diacetate into its non-fluorescent form, DAF-FM. In the presence of NO, DAF-FM is converted to a highly fluorescent triazole derivative. Fluorescence images were obtained using an inverted fluorescence microscope (Nikon, Tokyo, Japan).

### In Vitro Tube Formation Assay

Basement membrane matrix (Matrigel®, Corning, New York, USA) was added to a 4-well chamber slide and allowed to polymerize at 37°C for 30 min. Human umbilical vein endothelial cells (HUVECs; 2 × 10⁵) were mixed with hiPSC-derived endothelial cells (hiPSC-ECs; 2 × 10⁴) labeled with CellTracker™ CM-DiI (Invitrogen). The mixed cells were plated onto Matrigel-coated wells in EGM-2 medium (Lonza) and incubated at 37°C for 16–24 h. After incubation, cells were fixed with 4% paraformaldehyde (PFA) and counterstained with DAPI (Invitrogen). Tube-like structures were visualized using a fluorescence microscope (Nikon), and quantitative analysis of tube length, branch number, and tube number was performed using ImageJ software (NIH).

### Transplantation of Cells into the Ischemic Hindlimb

Hindlimb ischemia (HLI) surgery and cell transplantation were performed as previously described.^26^ In brief, 8- to 12-week-old male athymic nude mice (BALB/c-nu) underwent ligation of the femoral artery, and major collateral branches were cauterized to induce ischemia. Mice were randomly assigned to one of three groups: phosphate-buffered saline (PBS) only, non-PAD hiPSC-ECs, or PAD hiPSC-ECs. In all experimental groups, 2 × 10⁵ cells were injected into the ischemic hindlimb muscles immediately following surgery. For histological tracking, cells were pre-labeled with CellTracker™ CM-DiI (Invitrogen) before transplantation. Throughout the study, animals were maintained under specific pathogen–free conditions with sterile food and water, housed at a density of up to five mice per cage, and monitored according to Institutional Animal Care and Use Committee (IACUC) guidelines. Predefined exclusion criteria included systemic illness, respiratory distress, toxicity, inability to eat or drink, and body weight loss exceeding 15%. All animals that remained healthy during the observation period were included in the analyses.

### Measurement of Blood Flow in the Ischemic Hindlimb

Blood flow in the ischemic hindlimb was assessed using a laser Doppler perfusion imager (Moor Instruments Ltd., Devon, UK) before surgery, immediately after surgery, and weekly for 4 weeks.^48^ Perfusion was quantified from stored digital, color-coded images, and mean perfusion values were calculated for each limb. To account for variations caused by ambient light or temperature, blood flow in the ischemic (left) limb was normalized to that of the contralateral, non-ischemic (right) limb.

### Scoring for Limb Loss

Hindlimb necrosis was evaluated four weeks after surgery. Limb loss severity was scored on a six-point scale as follows: 0 = none; 1 = tip necrosis; 2 = toe necrosis; 3 = foot necrosis; 4 = leg necrosis; and 5 = complete limb loss.^59^

### Measurement of Capillary Density

To assess capillary density, mice with or without hindlimb ischemia were systemically perfused with FITC-conjugated isolectin B4 (ILB4; Vector Laboratories) immediately before euthanasia. Harvested tissues were fixed, sectioned, and imaged using a Zeiss LSM 700/780 confocal laser scanning microscope equipped with LSM Image software (Carl Zeiss). Quantitative analysis of capillary density was performed from multiple randomly selected microscopic fields.

### Histologic Analysis

Before euthanasia, mice were intravenously injected with fluorescein-conjugated *Griffonia (Bandeiraea) simplicifolia* isolectin B4 (ILB4; Vector Laboratories, Burlingame, USA) as previously described.^59^ Fifteen minutes after injection, mice were euthanized and perfused with normal saline to remove blood cells and unbound ILB4, followed by fixation with 4% paraformaldehyde (PFA). Hindlimb tissues were dissected, fixed in 4% PFA at 4°C for 16 h, and cryoprotected in 30% sucrose for 24 h. Frozen tissue sections (10–50 μm thick) were prepared using a cryostat. Images of engrafted cells and vascular structures were obtained by an investigator blinded to group allocation using a Zeiss LSM 700/780 Meta confocal laser scanning microscope equipped with a 20× objective and ZEN Black software (Carl Zeiss).

### Statistical Analysis

Unless otherwise indicated, data are presented as mean ± SEM. RNA-seq box plots show the median and interquartile range, with whiskers extending to 1.5 times the interquartile range. Donor age and BMI were compared between the non-PAD and PAD groups using two-sided Mann–Whitney U tests, and categorical demographic and clinical variables were compared using two-sided Fisher’s exact tests. Statistical significance for other experiments was determined using unpaired t tests, one-way ANOVA with Tukey’s multiple-comparison test, or two-way ANOVA with the two-stage step-up method, as appropriate. Analyses were performed using GraphPad Prism 9 (GraphPad Software, San Diego, CA, USA), and P < 0.05 was considered statistically significant.

## RESULTS

### Demographic Characteristics of Donors

We generated hiPSCs from eight PAD patients and seven non-PAD volunteers. This study included volunteers over 50 years old, as PAD typically occurs in individuals aged over 50.^60, 61^ For the PAD group, we only included patients with Rutherford categories 4-5.^62–64^ For the non-PAD group, we included volunteers with no clinical cardiovascular diseases or genetic diseases (Table S1). Demographic and clinical characteristics showed no significant differences in age, gender, hypercholesterolemia, body mass index (BMI), myocardial infarction (MI), end-stage renal disease (ESRD), previous amputation, hypertension, or chronic renal failure (CRF) between the two groups. The prevalence of coronary artery disease (CAD) and diabetes was significantly higher in PAD patients than in non-PAD donors (Figure S1 and Table S1).

### Generation and Characterization of hiPSCs Derived from non-PAD and PAD Donors

We generated hiPSCs using episomal vectors from peripheral blood through three stages (Figure S2A): erythroblast enrichment, electroporation, and iPSC initiation, as previously described.^47, 65^ First, peripheral blood mononuclear cells (PBMCs) were isolated from the donor’s blood, and erythroblasts were expanded from PBMCs. Erythroblasts were then electroporated using episomal vectors (Epi5^TM^ Episomal iPSC Reprogramming Kit, Thermo Fisher Scientific). Seven to twenty-one days after electroporation (mean, 16 days), hiPSC colonies were isolated, and the feeder-free transfer method was used for passaging the cells. Typically, 4-5 passages (∼60 days) were required to establish the cell lines. Once the hiPSCs were established, around passages 8–15, through the formation of hESC-like round colonies consisting of 200 to 300 cells, the stemness of the hiPSCs was verified. qRT-PCR, immunostaining, flow cytometry, chromosomal analyses, and a teratoma assay were performed. A human ESC line, H9, and a qualified hiPSC line, BJ1, were used as positive controls.^66^ qRT-PCR analysis of seven pluripotency genes, *OCT4, NANOG, SOX2, REX1, ESG1, DNMT3B*, and *GDF3*, showed that the mRNA levels were consistent between the non-PAD and PAD groups and were comparable to those of H9-ESCs and BJ1-hiPSCs (Figure 1A). When the data were analyzed by individual cell lines, statistical differences were observed, but no differences were noted among the categorized groups (Figure S3). Immunostaining demonstrated that all hiPSC lines expressed embryonic stem cell markers such as OCT4, SOX2, SSEA4, and TRA-1-60 (Figure 1B). Flow cytometry for pluripotency markers, TRA-1-60 and SSEA4, also showed no difference between the non-PAD and PAD groups (Figure 1C). We conducted karyotyping to evaluate genetic stability during hiPSC generation. Chromosomal analysis of all hiPSC lines showed a normal karyotype (Figure S4A). Furthermore, to assess the pluripotency of the hiPSCs in vivo, we performed a teratoma formation assay (Figure S4B). We subcutaneously injected hiPSCs (1 x 10^7^) into immunodeficient mice. After 6-12 weeks, we observed well-encapsulated cystic tumors in all mice, which contained differentiated elements from all three primary embryonic germ layers: endoderm (gut epithelium), mesoderm (cartilage), and ectoderm (neural tissue), indicating teratoma formation. Together, these data demonstrate that all the generated hiPSCs exhibited pluripotency and genetic stability. Furthermore, there were no significant differences in the hiPSC lines between the non-PAD and PAD groups.

**Figure 1.**
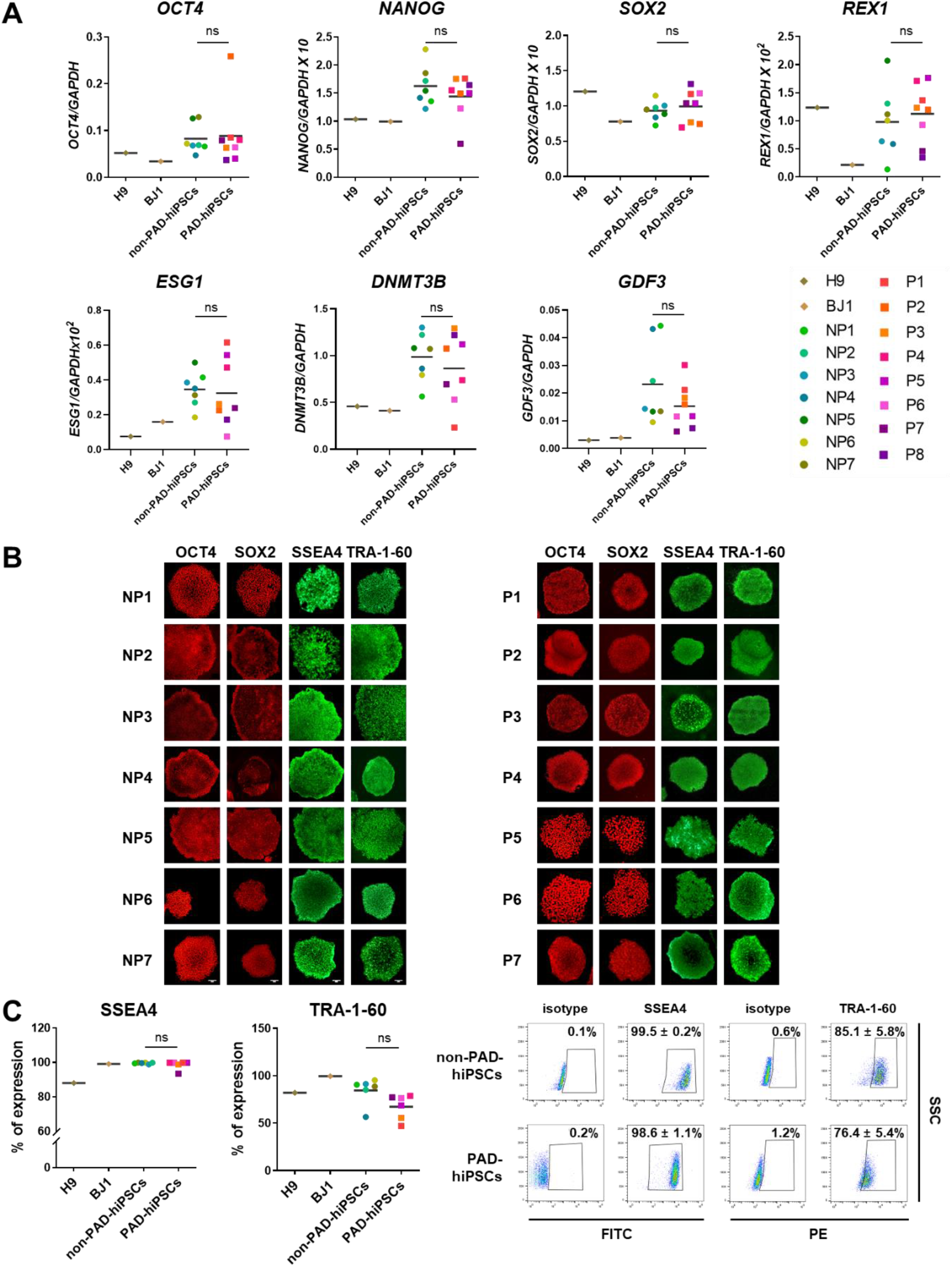
Characterization of Stemness in Generated hiPSCs. **A**, Quantitative RT-PCR analysis of pluripotency-related genes in hiPSCs. Relative mRNA expression was normalized to *GAPDH* (set to 1). Data represent three independent experiments with technical triplicates for each cell line (n = 7 for non-PAD; n = 8 for PAD). Statistical analysis was performed using one-way ANOVA followed by Tukey’s multiple comparison test. Data are shown as mean ± SEM. ns, not significant. **B**, Immunostaining of hiPSCs for pluripotency markers OCT4, SOX2, SSEA4, and TRA-1-60. **C**, Flow-cytometric analysis of SSEA4 and TRA-1-60 expression in hiPSCs. IgG isotype antibodies served as negative controls. Statistical analysis was conducted using an unpaired t-test. Data are shown as mean ± SEM (n = 7 for non-PAD; n = 8 for PAD).

### Transcriptomic Expression Profiles of hiPSCs Derived from non-PAD and PAD Donors

To gain more detailed insight into the transcriptomic profiles of hiPSCs from non-PAD and PAD donors, we performed genome-wide RNA sequencing (RNA-seq) analysis. Principal component analysis (PCA) showed overlapping distributions among hiPSC samples from both groups and no distinct separation in global gene expression (PC1 with 21.6% variance and PC2 with 16% variance, Figure 2A). Hierarchical clustering and heatmap analysis based on the top 1,000 most variable genes across samples showed no clear clustering according to donor group (Figure 2B). Scatter plot analysis showed that gene expression profiles remained largely consistent between the non-PAD and PAD groups in hiPSCs (Figure 2C). Transcriptomic analysis identified 262 differentially expressed genes (DEGs) among 16,337 genes tested (1.6%), including 196 genes upregulated and 66 genes downregulated in PAD hiPSCs compared with non-PAD hiPSCs. We then analyzed 82 pluripotency-related genes (Table S3) and found that hiPSCs from non-PAD and PAD donors exhibited equivalent expression patterns for these genes (Figure 2D and Figure S5A and B), consistent with the qRT-PCR results. Focused analysis of 82 predefined pluripotency-related genes identified eight genes (9.8%) with nominal differences between non-PAD and PAD hiPSCs at the raw p-value level (*DDX21, DPPA2, DPPA5, ERAS, GDF3, KHDC3L, MFGE8*, and *PARP1*). Among these, *DDX21, DPPA2, DPPA5, ERAS, GDF3, KHDC3L*, and *PARP1* showed lower expression in PAD hiPSCs, whereas *MFGE8* was increased. However, after Benjamini–Hochberg correction for multiple testing, only *KHDC3L* remained significant (1.2%). Its expression level was low in both groups, but was reduced in PAD hiPSCs compared with non-PAD hiPSCs. Given that *KHDC3L* is primarily associated with oocyte and early embryonic programs rather than the core pluripotency network, this isolated difference is more appropriately interpreted as limited gene-level variation than as evidence of altered pluripotent identity. Overall, these findings indicate that PAD status does not broadly perturb the pluripotency-associated transcriptional landscape of hiPSCs.

**Figure 2.**
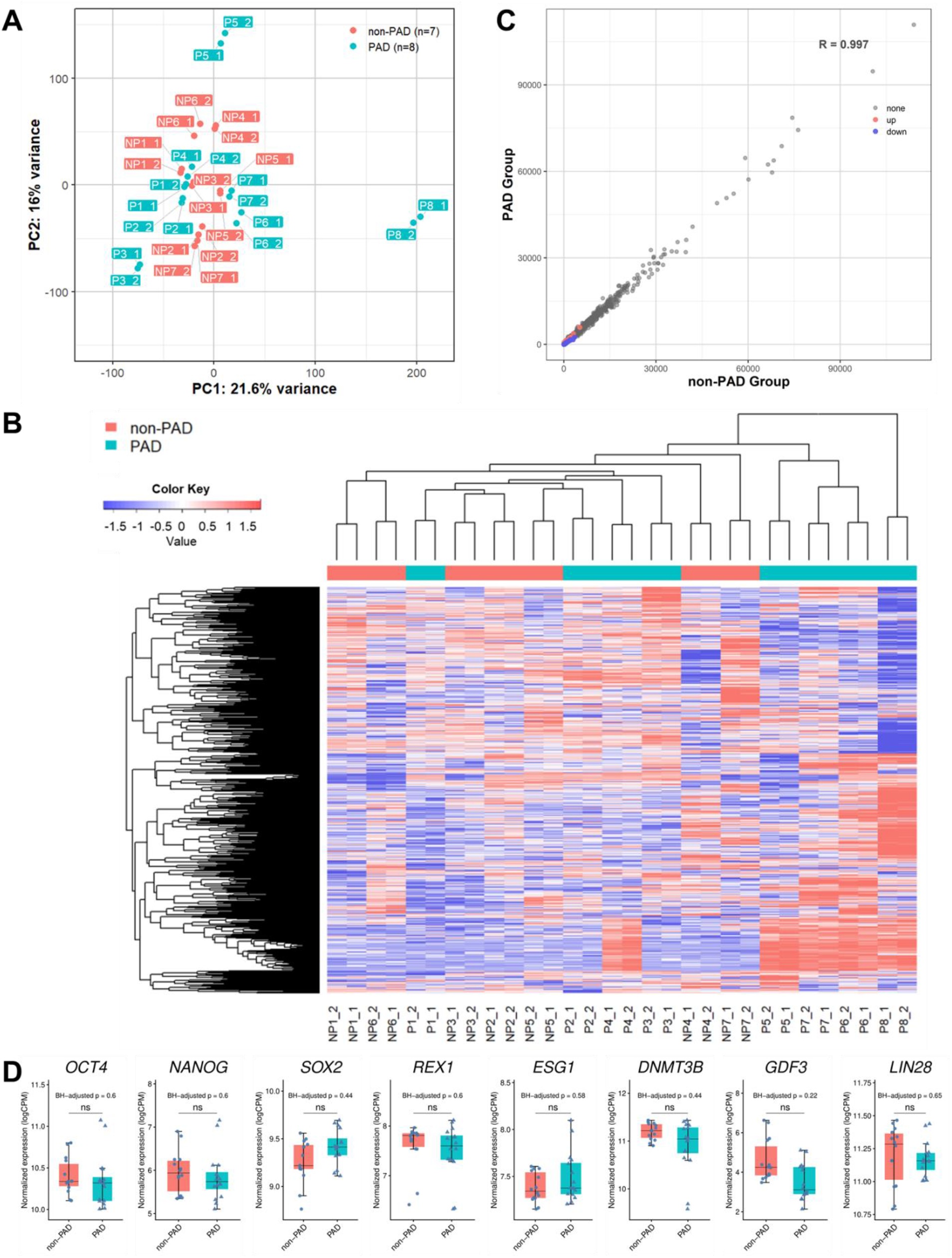
Transcriptomic Analysis of hiPSCs by RNA Sequencing. **A**, PCA of hiPSCs derived from non-PAD (pink) and PAD (turquoise) donors. **B**, Hierarchical clustering heatmap of the top 1,000 most variable genes across hiPSC samples derived from non-PAD and PAD donors. Gene expression values are shown as row-scaled log2 counts per million (logCPM). The color scale ranges from blue (low relative expression) to red (high relative expression). Color annotations indicate non-PAD (pink) and PAD (turquoise) samples. **C**, Scatter plot showing Pearson correlation coefficients among hiPSC samples derived from non-PAD and PAD donors. **D**, Box plots showing TMM-normalized log2 counts per million (logCPM) values for eight representative pluripotency-related genes in hiPSCs derived from non-PAD and PAD donors. Each point represents one replicate sample (14 non-PAD samples from 7 donor-derived lines and 16 PAD samples from 8 donor-derived lines; two replicate samples per line). Boxes indicate the median and interquartile range, and whiskers extend to 1.5 times the interquartile range. P values were calculated using the two-sided Wilcoxon rank-sum test and adjusted for multiple comparisons using the Benjamini–Hochberg method. BH-adjusted p values are shown. ns, not significant.

### Characterization of hiPSC-ECs Derived from non-PAD and PAD Donors

Next, we compared the characteristics of hiPSC-ECs between the non-PAD and PAD groups. hiPSCs were differentiated into ECs (hiPSC-ECs) using the protocol established previously.^26^ Briefly, hiPSCs were first induced into the mesodermal lineage for three days. These mesodermally differentiated hiPSCs were then cultured in medium containing VEGFA and FGF2 for an additional 11 to 13 days. The resulting hiPSC-ECs were replated for expansion and maintained through serial passaging (Figure S2B). Cells were characterized at passage 2. Gene expression analysis of representative EC markers such as *CDH5, PECAM1, KDR, VWF, TEK,* and *NOS3* by qRT-PCR confirmed their expression in hiPSC-ECs differentiated from all hiPSC lines, with no significant difference between the non-PAD and PAD groups (Figure 3A). Flow cytometric analysis of hiPSC-ECs also demonstrated expression of CDH5 (∼94%), PECAM1 (∼65%), VWF (∼72%), and KDR (∼84%), with no significant difference in marker expression between the two groups (Figure 3B). Immunostaining further confirmed expression of CDH5 and KDR in all cell lines (Figure 3C). Collectively, these data indicate that the generated hiPSC-ECs exhibited endothelial lineage characteristics, and there were no statistically significant differences in endothelial features between the non-PAD and PAD groups.

**Figure 3.**
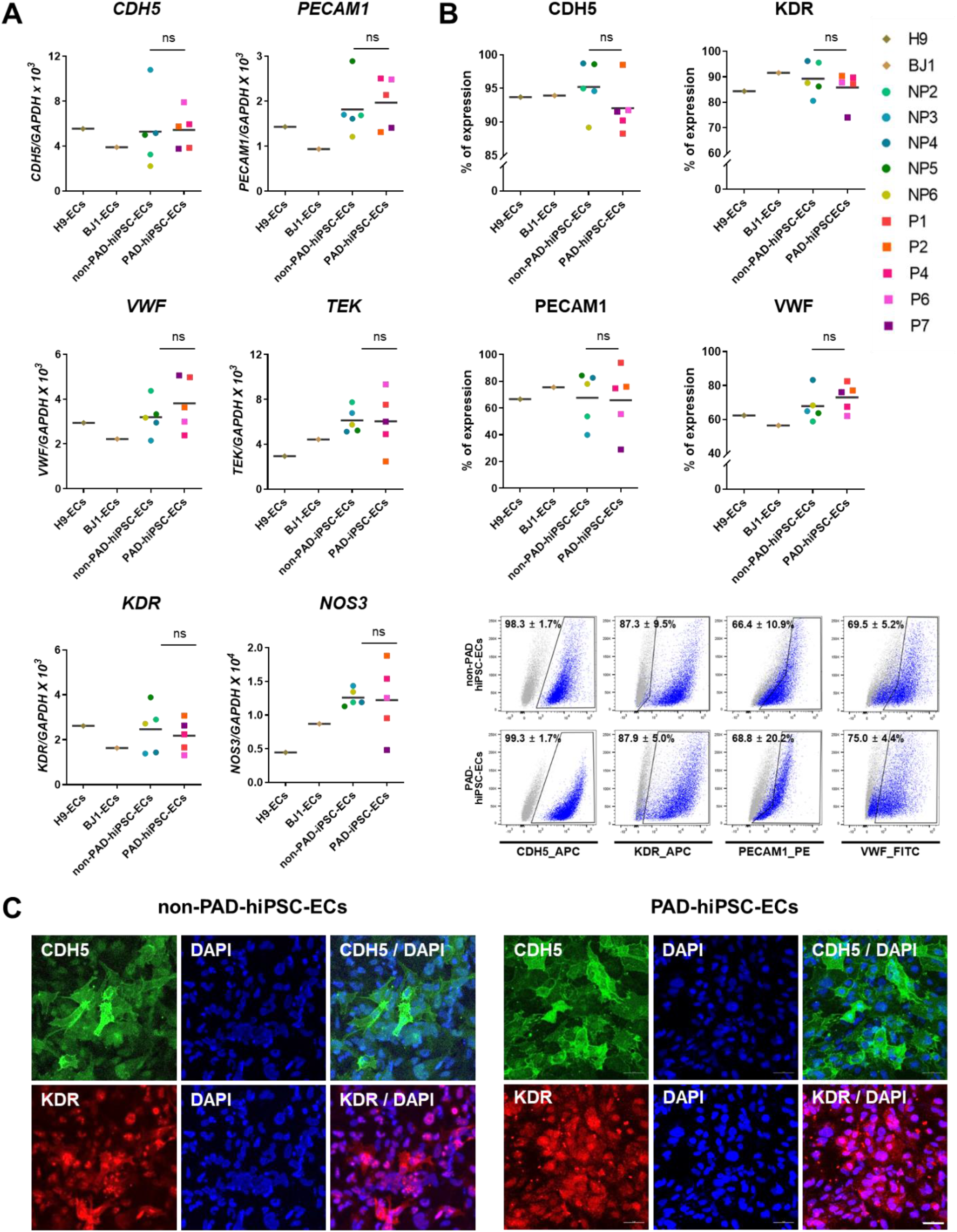
Endothelial Characterization of hiPSC-ECs. **A**, Quantitative RT-PCR analysis of endothelial cell–specific genes in hiPSC-ECs. Relative mRNA expression was normalized to *GAPDH* (set to 1). Data represent three independent experiments with technical triplicates for each cell line (n = 5 for non-PAD; n = 5 for PAD). Statistical analysis was performed using one-way ANOVA followed by Tukey’s multiple comparison test. Data are shown as mean ± SEM. ns, not significant. **B**, Flow-cytometric analysis of CDH5, KDR, PECAM1, and VWF expression in hiPSC-ECs. IgG isotype antibodies served as negative controls. Statistical analysis was performed using one-way ANOVA with Tukey’s multiple comparison test. Data are shown as mean ± SEM (n = 5 for non-PAD; n = 5 for PAD). ns, not significant. **C**, Immunostaining of hiPSC-ECs for endothelial markers CDH5 and KDR.

### Transcriptomic Profiles of hiPSC-ECs Derived from non-PAD and PAD Donors

To compare the global transcriptomic landscapes of hiPSC-ECs from non-PAD and PAD donors, we performed RNA-seq. PCA showed overlapping distributions among hiPSC-EC samples from both groups, with no distinct separation in global gene expression (PC1, 26.9% variance; PC2, 14.5% variance; Figure 4A). Hierarchical clustering and heatmap analysis based on the top 1,000 most variable genes across samples showed no clear segregation according to donor group (Figure 4B). Scatter plot analysis further indicated that gene expression profiles were largely preserved between the non-PAD and PAD groups in hiPSC-ECs (Figure 4C). Transcriptomic analysis identified 51 differentially expressed genes (DEGs) among 15,290 genes tested (0.3%), including 21 genes upregulated and 30 genes downregulated in PAD hiPSC-ECs compared with non-PAD hiPSC-ECs. We next focused on 90 endothelial cell–related genes (Table S4) and found that hiPSC-ECs from non-PAD and PAD donors exhibited equivalent expression patterns of these genes (Figure 4D and Figure S6A and B), consistent with our qRT-PCR data. Focused analysis of 90 predefined EC-related genes identified ten genes (11.1%) with nominal differences between non-PAD and PAD hiPSC-ECs at the raw p-value level (*ACVRL1, ANXA5, CD34, EGFL7, EPOR, FABP5, IL13RA1, NOS3, RIPK1*, and *S1PR1*). Among these, *ACVRL1, CD34, EGFL7, EPOR, IL13RA1, NOS3, RIPK1*, and *S1PR1* showed lower expression in PAD hiPSC-ECs, whereas *ANXA5* and *FABP5* were increased. However, after Benjamini–Hochberg correction for multiple testing, none of these EC-related genes remained significant (0%). Overall, these findings indicate that PAD status does not broadly perturb the EC-associated transcriptional landscape of hiPSC-ECs.

**Figure 4.**
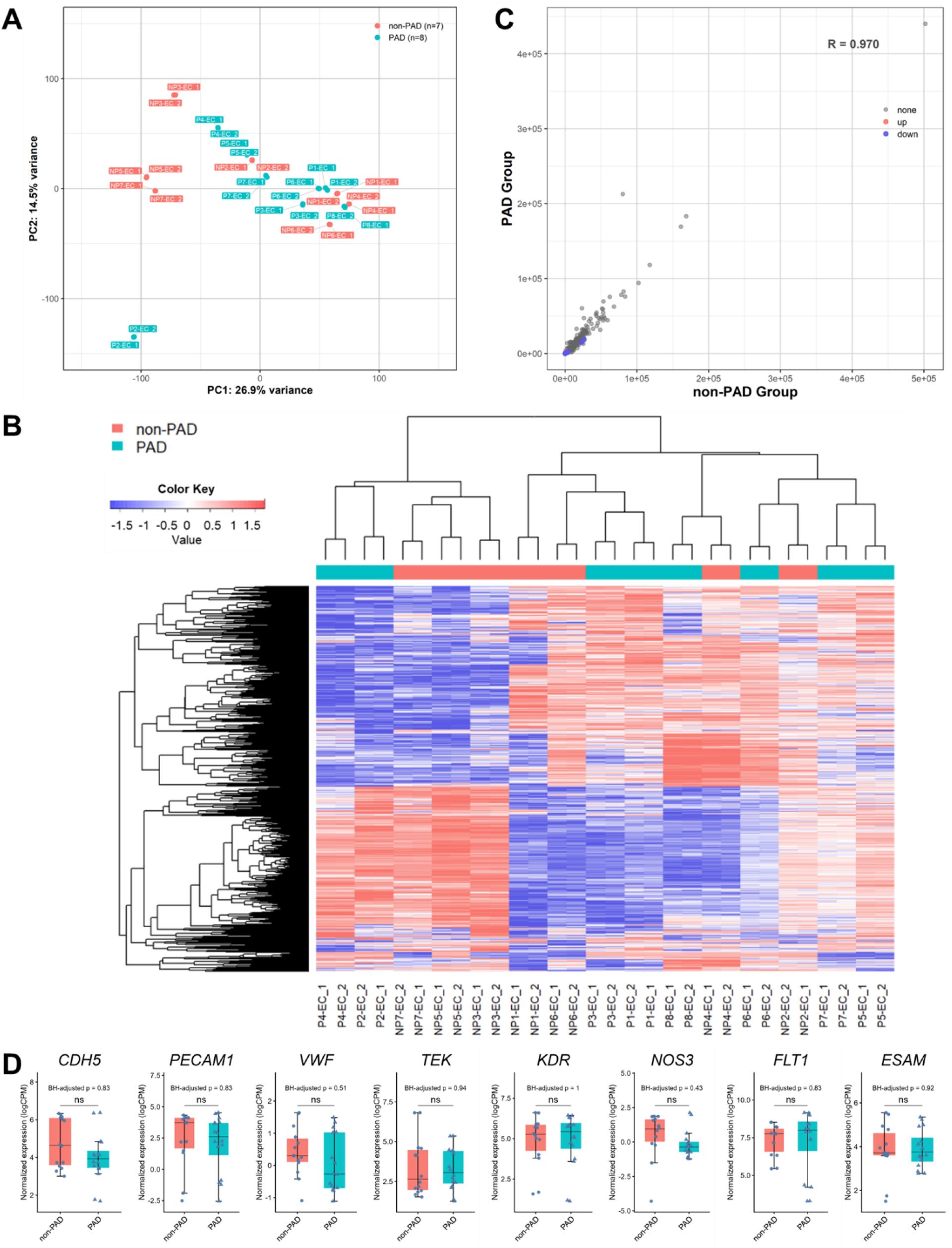
Transcriptomic Analysis of hiPSC-ECs by RNA Sequencing. **A**, PCA of hiPSC-ECs derived from non-PAD (pink) and PAD (turquoise) donors. **B**, Hierarchical clustering heatmap of the top 1,000 most variable genes across hiPSC-EC samples derived from non-PAD and PAD donors. Gene expression values are shown as row-scaled log2 counts per million (logCPM). The color scale ranges from blue (low relative expression) to red (high relative expression). Color annotations indicate non-PAD (pink) and PAD (turquoise) samples. **C**, Scatter plot showing Pearson correlation coefficients among hiPSC-EC samples derived from non-PAD and PAD donors. **D**, Box plots showing TMM-normalized log2 counts per million (logCPM) values for eight representative endothelial cell–related genes in hiPSC-ECs derived from non-PAD and PAD donors. Each point represents one replicate sample (14 non-PAD samples from 7 donor-derived lines and 16 PAD samples from 8 donor-derived lines; two replicate samples per line). Boxes indicate the median and interquartile range, and whiskers extend to 1.5 times the interquartile range. P values were calculated using the two-sided Wilcoxon rank-sum test and adjusted for multiple comparisons using the Benjamini– Hochberg method. BH-adjusted p values are shown. ns, not significant.

### Comparison of Transcriptomic Profiles between hiPSCs and hiPSC-ECs

To investigate transcriptomic alterations associated with endothelial differentiation, RNA sequencing data from hiPSCs and hiPSC-ECs derived from non-PAD and PAD donors were comparatively analyzed. PCA demonstrated a clear separation between hiPSCs and hiPSC-ECs, indicating that global transcriptomic profiles were primarily determined by cell identity rather than donor disease status (Figure S7A). Hierarchical clustering analysis further confirmed this distinction, as samples clustered predominantly according to cell type irrespective of PAD status (Figure S7B). Consistently, Pearson correlation analysis revealed strong positive correlations among samples within the same cell type, whereas correlations between hiPSCs and hiPSC-ECs were comparatively lower, reflecting substantial transcriptional reprogramming during endothelial differentiation (Figure S7C). K-means clustering analysis of the top 3,000 differentially expressed genes identified two major gene clusters associated with cell identity (Figure S7D). Cluster A consisted of 1,298 genes preferentially expressed in hiPSCs, whereas Cluster B consisted of 1,702 genes enriched in hiPSC-ECs. Gene Ontology (GO) Biological Process enrichment analysis demonstrated that genes enriched in the hiPSC cluster were primarily associated with developmental and morphogenetic processes, including stem cell differentiation, embryonic morphogenesis, and cell fate commitment. In contrast, genes enriched in the hiPSC-EC cluster were predominantly associated with vascular and endothelial functions, including blood vessel development, vasculature development, angiogenesis, and endothelial cell migration, supporting successful endothelial lineage specification following differentiation (Figure S7E). Together, these results indicate that differentiation from hiPSCs to hiPSC-ECs induces extensive transcriptomic remodeling, whereas donor disease status (PAD versus non-PAD) exerts relatively limited effects on global gene expression patterns within each cell type.

### In Vitro Functionality of hiPSC-ECs

To assess the in vitro functionality of hiPSC-ECs, nitric oxide (NO) production was measured.^26^ hiPSC-ECs derived from both non-PAD and PAD donors exhibited the characteristic cobblestone-like morphology of endothelial cells and produced NO, as detected by DAF-FM diacetate staining. DAF-FM fluorescence intensity did not differ significantly between the non-PAD and PAD groups (Figure 5A). Next, to evaluate vessel-forming capacity, a modified Matrigel tube formation assay was performed by co-culturing HUVECs and hiPSC-ECs. Prelabeled hiPSC-ECs with the red fluorescent dye DiI integrated into tubular networks with HUVECs. Quantitative analyses of branch number, tube number, and total tube length showed no significant differences between the non-PAD and PAD groups (Figure 5B). Together, these findings indicate that hiPSC-ECs derived from both donor types maintain endothelial functionality, including NO production and tube-forming ability, regardless of PAD status.

**Figure 5.**
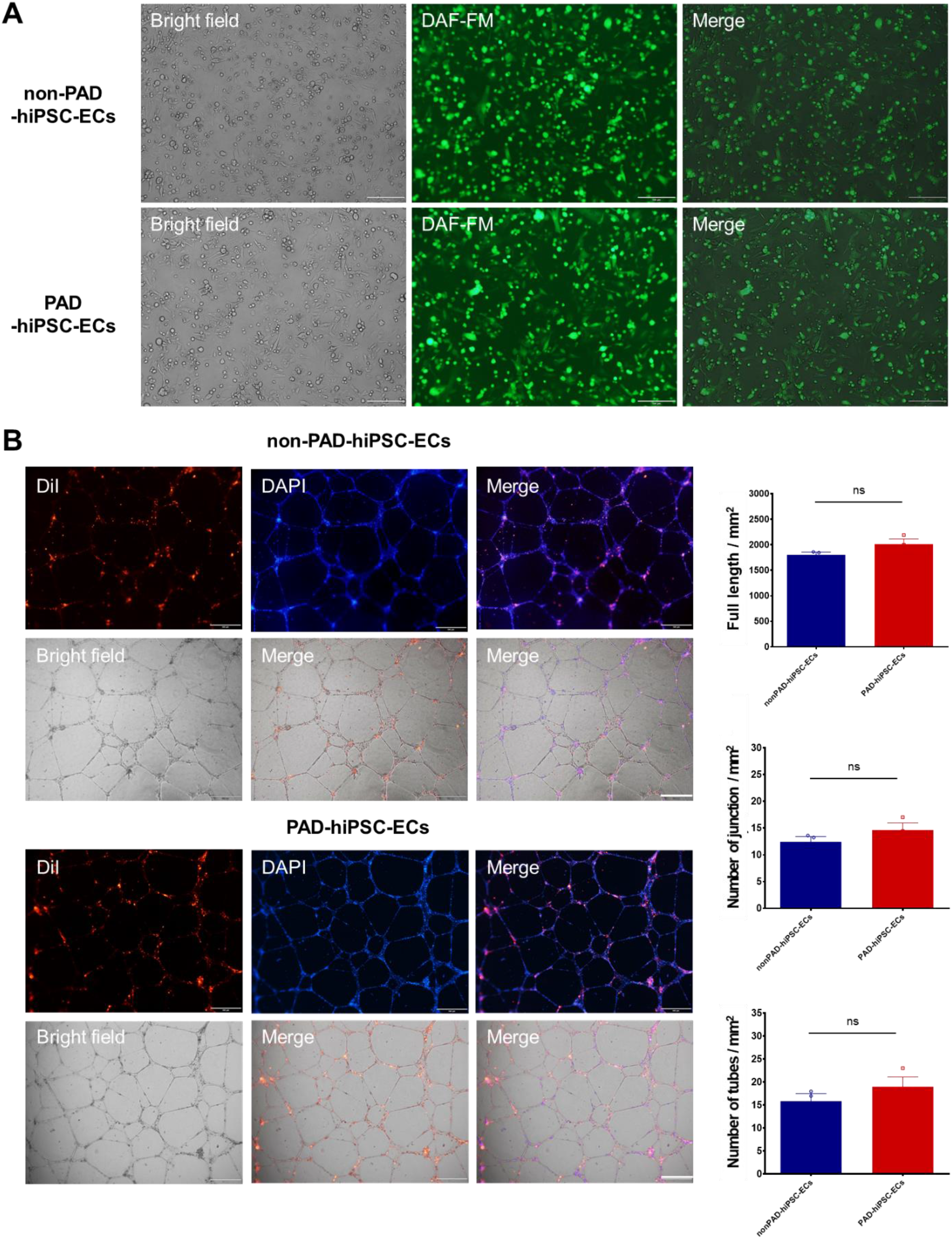
In Vitro Functional Assessment of hiPSC-ECs. **A**, Representative phase-contrast and fluorescence images of hiPSC-ECs after incubation with DAF-FM diacetate, a fluorescent indicator of nitric oxide (NO) production (green). **B**, Representative phase-contrast and fluorescence images of tube formation on Matrigel after co-culture of DiI-labeled hiPSC-ECs (red) with HUVECs. Nuclei are stained with DAPI (blue). Quantification of tube formation (right) shows the number of branches, number of tubes, and total tube length. Data represent three independent experiments, with three randomly selected fields quantified per cell line. Statistical analysis was performed using one-way ANOVA followed by Tukey’s multiple comparison test. Data are shown as mean ± SEM (n = 3 for non-PAD; n = 3 for PAD).

### Therapeutic and Vessel-Forming Effects of hiPSC-ECs in Hindlimb Ischemia

To evaluate the therapeutic potential of hiPSC-ECs, cells were transplanted into the ischemic hindlimb muscle of mice following femoral vessel ligation. Hindlimb ischemia (HLI) was surgically induced in nude mice, and hiPSC-ECs (2 × 10⁵; three lines each from non-PAD and PAD groups) or PBS were injected into the ischemic muscle (Figure S2C).^26^ Laser Doppler perfusion imaging revealed that hiPSC-EC transplantation markedly enhanced blood flow recovery at days 7, 14, 21, and 28 compared with PBS treatment, with no significant difference between the non-PAD and PAD groups (Figure 6A). Limb loss scores were significantly improved in both hiPSC-EC–treated groups compared with PBS, whereas no difference was detected between the non-PAD and PAD groups (Figure 6B). Capillary density analysis in tissues harvested at day 28, after systemic perfusion with FITC-conjugated ILB4, showed higher vessel density in both transplantation groups than in PBS controls, with no discernible difference between donor origins (Figure 6C). Collectively, these results demonstrate that hiPSC-ECs from both non-PAD and PAD donors effectively promoted perfusion recovery, limb preservation, and neovascularization in ischemic tissue, indicating comparable therapeutic efficacy and vessel-forming capacity regardless of PAD status.

**Figure 6.**
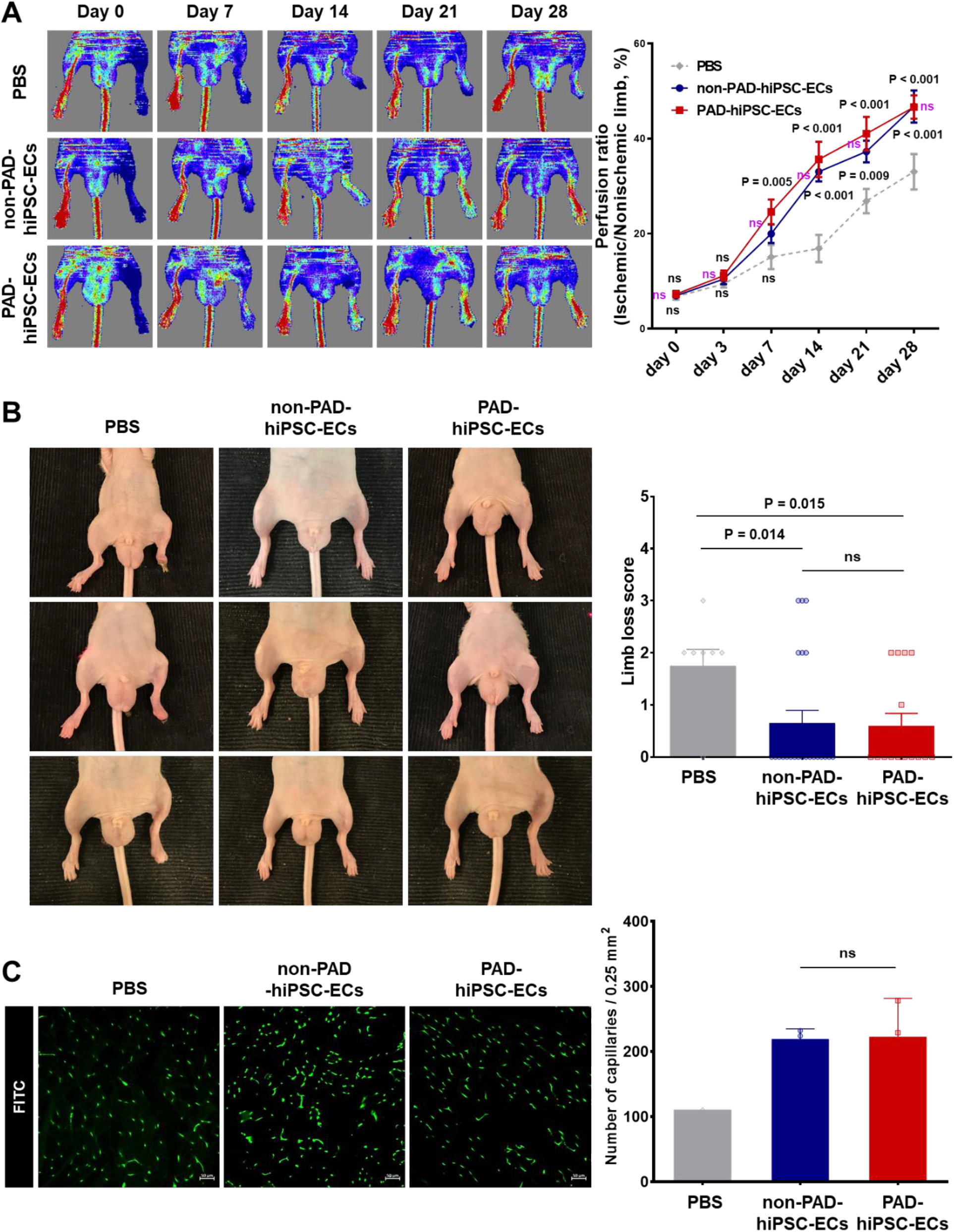
Therapeutic Effects of hiPSC-ECs in a Mouse Hindlimb Ischemia Model. **A**, Representative laser Doppler perfusion images (left) and quantitative analysis (right) of mice treated with PBS, non-PAD hiPSC-ECs, or PAD hiPSC-ECs over a 28-day period. Statistical analysis was performed using two-way ANOVA followed by the two-stage step-up method (n = 7-29 per group). **B**, Representative photographs of mouse hindlimbs (top) and limb-loss score index (bottom) at 28 days after implantation of PBS, non-PAD hiPSC-ECs, or PAD hiPSC-ECs. Statistical analysis was conducted using one-way ANOVA with Tukey’s multiple comparison test (n = 8-23 per group). **C**, Representative images of FITC-isolectin B4 (ILB4)–perfused vessels (top) and quantification of vascular density (bottom) at 28 days after implantation of PBS, non-PAD hiPSC-ECs, or PAD hiPSC-ECs. Statistical analysis was performed using one-way ANOVA with Tukey’s multiple comparison test (n = 10-18 per group).

### In Vivo Neovascularization of Engrafted hiPSC-ECs in Hindlimb Ischemia

We next assessed the vasculogenic activity and direct vessel-forming contribution of engrafted hiPSC-ECs through histological analyses. To visualize functional endothelium, FITC-conjugated ILB4 was injected into the heart before sacrifice. Confocal microscopy revealed that engrafted hiPSC-ECs from both non-PAD and PAD donors were frequently localized near host vasculature and integrated into functional vessels. Orthogonal confocal imaging further confirmed the incorporation of hiPSC-ECs into host vessels in both groups (Figure 7A, Figure 7B, and Videos S1, S2). Notably, subsets of engrafted hiPSC-ECs in both non-PAD and PAD groups exhibited incorporation into the vascular wall, perivascular localization, and a guiding pattern, aligned linearly along vascular structures. The incorporation, perivascular localization, and guiding alignment of engrafted hiPSC-ECs from PAD donors were comparable to those observed in cells derived from non-PAD donors (Figure 7A, Figure 7B, and Videos S1, S2). Together, these findings indicate that engrafted hiPSC-ECs from both donor types possess equivalent angiogenic and vessel-forming capabilities in vivo.

**Figure 7.**
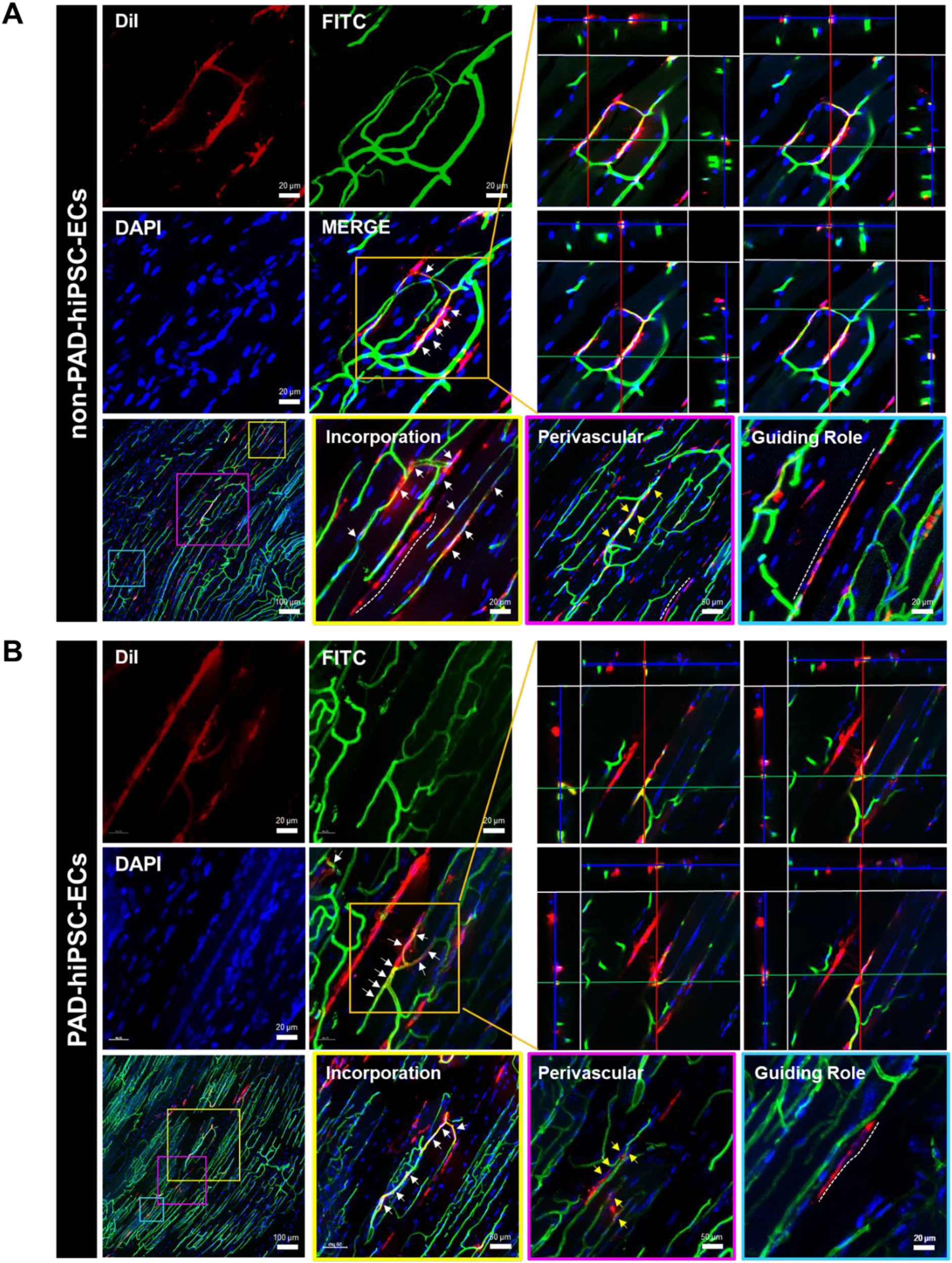
Contribution of Engrafted hiPSC-ECs to Neovascularization. **A** and **B,** Representative confocal images illustrating the localization patterns of transplanted non-PAD hiPSC-ECs (**A**) and PAD hiPSC-ECs (**B**) in ischemic hindlimb tissue 28 days after implantation. Transplanted hiPSC-ECs were identified by DiI labeling (red), host vasculature was visualized by FITC perfusion (green), and nuclei were counterstained with DAPI (blue). Representative images of vascular incorporation, perivascular localization, and guiding roles during neovascularization are shown. White arrows indicate incorporated hiPSC-ECs within vascular structures, yellow arrows indicate perivascularly localized hiPSC-ECs, and white dotted lines indicate guiding roles during vessel formation. Upper right panels show orthogonal confocal images from each group. Lower panels show representative high-magnification images of vascular incorporation, perivascular localization, and guiding roles in the non-PAD and PAD groups.

## DISCUSSION

In this study, we generated hiPSCs from the peripheral blood of seven non-PAD donors and eight PAD patients and conducted a comprehensive, side-by-side comparison of their molecular, cellular, and functional characteristics. hiPSCs from both groups expressed pluripotency markers, maintained normal karyotypes, and formed teratomas containing derivatives of all three embryonic germ layers. Genome-wide RNA-seq revealed comparable global transcriptomic profiles. Upon directed differentiation using a clinically compatible protocol, hiPSC-ECs from non-PAD and PAD donors displayed equivalent expression of endothelial markers at both the transcript and protein levels, with no major differences in global gene expression. Functionally, hiPSC-ECs from both groups exhibited preserved nitric oxide production and tube-forming capacity in vitro. In a murine hindlimb ischemia model, hiPSC-ECs from either source exhibited comparable therapeutic potency, as evidenced by perfusion recovery, limb salvage, and neovascularization, and engrafted cells integrated into host vasculature with similar efficiency. To our knowledge, this is the first large-scale, head-to-head comparison of hiPSCs and hiPSC-ECs derived from PAD patients and non-PAD donors that integrates molecular, cellular, and in vivo functional assessments.

At the hiPSC stage, no consistent molecular or cellular differences were detected between the non-PAD and PAD groups. Although individual cell lines exhibited statistically significant variability in pluripotency-gene expression by qRT-PCR, no systematic group-level differences emerged. This pattern is most plausibly explained by inter-individual variation rather than disease-specific effects. Each hiPSC line displays intrinsic heterogeneity in morphology, growth dynamics, gene expression, and lineage propensity, and because hiPSC generation and differentiation involve multiple sequential steps, minor variations can accumulate to produce distinct line-to-line outcomes. Such heterogeneity remains a recognized challenge for hiPSC-based therapeutics.^67^

Epigenetic memory likely contributes to this variability. Prior studies have shown that hiPSCs retain residual epigenetic features inherited from their parental cells,^68–70^ and that donor cell type influences the epigenome and differentiation potential of derived hiPSCs. Donor-level genetic variation further shapes hiPSC phenotypes: the Human Induced Pluripotent Stem Cells Initiative (HipSci; http://www.hipsci.org) reported that 5–46% of variation in hiPSC phenotypes, including differentiation capacity and cellular morphology, can be attributed to inter-individual differences.^71^ Collectively, these data indicate that, in non-genetic diseases such as PAD, donor-specific variation likely outweighs disease-related effects, particularly because epigenetic signatures tend to diminish with continued passaging.^72^ We further speculate that hiPSCs and their derivatives share conserved molecular pathways during reprogramming and differentiation regardless of donor disease status. Consistent with this, although RNA-seq identified 262 DEGs between groups (1.6% of all detected genes), no major changes were observed in core pluripotency regulators or reprogramming factors. Among 82 pluripotency-related genes examined, only KHDC3L remained significant after Benjamini–Hochberg correction, and its expression was low in both groups. Given that KHDC3L is associated with oocyte and early embryonic programs rather than the core pluripotency network,^73–75^ this isolated finding is best interpreted as gene-level variation rather than evidence of altered pluripotent identity.

hiPSC-ECs derived from non-PAD and PAD donors likewise showed no significant molecular, cellular, or in vivo differences. Endothelial marker expression, in vitro functionality (NO production, tube formation), and in vivo therapeutic efficacy (perfusion recovery, limb salvage, neovascularization) were all comparable between groups. Engrafted hiPSC-ECs from both donor types integrated into host vasculature and displayed equivalent vessel-forming activity. As at the hiPSC stage, inter-line variability exceeded between-group differences, reinforcing the conclusion that donor-individual factors—rather than PAD status—drive the residual variation observed. These findings support the suitability of hiPSC-ECs for autologous, rather than allogeneic, cell therapy. RNA-seq identified modest transcriptional differences between groups at the hiPSC-EC stage, with only 0.3% of detected genes (51 of 15,290) classified as DEGs. Focused analysis of 90 EC-related genes identified ten genes with nominal differences at the raw p-value level, but none remained significant after Benjamini–Hochberg correction. Importantly, the number of DEGs between groups was lower in hiPSC-ECs than in hiPSCs, both for global and lineage-specific expression. This reduction suggests that endothelial differentiation following reprogramming progressively attenuates disease-associated transcriptional variation. Together, these observations indicate that reprogramming followed by lineage-directed differentiation largely resets disease-associated baseline gene expression, providing a mechanistic basis for the functional equivalence we observed between groups.

A central challenge in the clinical translation of hiPSC-based therapies is the potential tumorigenicity of hiPSCs and their derivatives.^67^ The virtually unlimited proliferative capacity of hiPSCs, an advantage for scalable manufacturing, also raises the risk of uncontrolled growth after transplantation. Three principal mechanisms warrant attention. First, residual undifferentiated or immature cells in the final product can give rise to teratomas; establishing fully defined, highly efficient differentiation protocols is therefore essential. The clinically compatible system^26^ used here yielded substantial enrichment of the endothelial population, with more than 98% of cells expressing CDH5. Second, residual reprogramming-factor activity may promote tumorigenesis. We used a non-integrating episomal vector system, which has been reported to be non-mutagenic during integration-free hiPSC induction.^76^ Previous work has shown that more than one-third of subclones lose episomal vectors by passages 9–10, and that episomal sequences become undetectable in most clones by passages 11–20.^65, 77^ In this study, hiPSCs at passages 15–25, at which episomal sequences were effectively absent were used for endothelial differentiation. Notably, the timing of vector loss varied even among clones from the same donor, underscoring the importance of considering both passage number and clonal variability during hiPSC generation. Third, tumorigenicity may arise from genetic mutations acquired during prolonged in vitro culture. In our histological analyses, no teratomas or tumors were observed in any mouse transplanted with hiPSC-ECs.

Our findings are consistent with, and extend, prior studies on patient-derived hiPSCs and their derivatives. hiPSCs from patients with monogenic disorders retain the causal genotype,^37–39^ and pathogenic mutations are preserved upon differentiation into relevant lineages.^29, 34, 40^ In contrast, hiPSCs from non-genetic diseases have generally shown characteristics comparable to those from healthy controls, although outcomes for differentiated derivatives have been mixed, with some studies reporting no differences and others reporting altered cellular phenotypes. Earlier studies were limited by small sample sizes or by reliance on in vitro assays alone. A recent study on PAD reported that hiPSC-ECs from a single PAD patient were phenotypically and functionally comparable to those from healthy donors, but was limited by an in vitro–only design and a single PAD donor.^44^ Notably, Song et al.^41^ recently described autologous hiPSC-derived midbrain dopaminergic progenitors transplanted into a patient with Parkinson’s disease, with sustained graft survival and symptomatic improvement at 24 months, providing landmark clinical proof of concept for autologous hiPSC-based therapy.^78^ Our study extends these prior reports by combining a clinically relevant cohort (eight PAD patients and seven non-PAD donors), a clinically compatible differentiation protocol, and integrated molecular, in vitro, and in vivo assessments.

Several limitations should be acknowledged. First, the PAD cohort included only Rutherford categories 4-5 patients; whether our findings generalize to more advanced disease (Category 6) or to patients with specific comorbid profiles warrants further investigation. Second, although hiPSC-ECs were assessed in a well-validated hindlimb ischemia model, long-term safety, tumorigenicity, and the durability of vascular regeneration beyond the observation window will need to be evaluated in additional preclinical studies before clinical application. Finally, although inter-individual variation appeared to dominate over disease-associated effects, the relative contributions of genetic background, epigenetic memory, and clonal heterogeneity to residual variability merit dedicated investigation.

In conclusion, this large-scale comparative study, employing a clinically compatible differentiation system, demonstrates that hiPSCs and hiPSC-ECs derived from PAD patients are molecularly, cellularly, and functionally equivalent to those derived from non-PAD donors, with comparable therapeutic potency in a hindlimb ischemia model. These findings establish that patient-derived hiPSCs and hiPSC-ECs provide a robust and clinically relevant platform for autologous cell therapy in ischemic vascular disease, supporting their further development as a personalized regenerative strategy for PAD.

## Supporting information

Video S1. Non-PAD hiPSC-EC neovascularization

Video S2. PAD hiPSC-EC neovascularization

## Nonstandard Abbreviations and Acronyms

BMI: body mass index
CAD: coronary artery disease
CLTI: chronic limb-threatening ischemia
CRF: chronic renal failure
CVD: cardiovascular disease
DAPI: 4′,6-diamidino-2-phenylindole
DAF-FM: 4-amino-5-methylamino-2′,7′-difluorofluorescein
DEG: differentially expressed gene
EC: endothelial cell
ESRD: end-stage renal disease
FDR: false discovery rate
FITC: fluorescein isothiocyanate
GO: Gene Ontology
GSEA: Gene Set Enrichment Analysis
hiPSC: human induced pluripotent stem cell
hiPSC-EC: human induced pluripotent stem cell-derived endothelial cell
HLI: hindlimb ischemia
HUVEC: human umbilical vein endothelial cell
ILB4: isolectin B4
NO: nitric oxide
PAD: peripheral artery disease
PBS: phosphate-buffered saline
PBMC: peripheral blood mononuclear cell
PCA: principal component analysis
PFA: paraformaldehyde
RNA-seq: RNA sequencing
qRT-PCR: quantitative real-time polymerase chain reaction

## Acknowledgements

Y.-s.Y. and S.-J.L. conceived the project and designed experiments. J.Y.B. performed most experiments. S.-J.L.2, Y.-G.K., and D.C. provided PBMCs of donors and technical advice. J.Y.B., H.-K.K., and H.O.K. generated hiPSCs. J.J. conducted qRT–PCR and flow cytometry analysis. S.P., J.J., and J.P. performed the hindlimb ischemia surgical experiments. J.K. and Y.K. carried out the teratoma formation assay. J.P. conducted immunostaining. J.Y.B., C.J., S.B., and J.W.H. contributed to RNA-seq data analysis. J.Y.B., S.-J.L., J.W.H., and Y.-s.Y. analyzed data, and J.Y.B., S.-J.L., and Y.-s.Y. wrote the manuscript.

## Supplemental Material

