## Supplemental Material for "Human iPSCs and iPSC-Derived Endothelial Cells from PAD Patients and Healthy Donors Exhibit Comparable Characteristics and Potency: Implications for Autologous Cell Therapy in Peripheral Artery Disease"

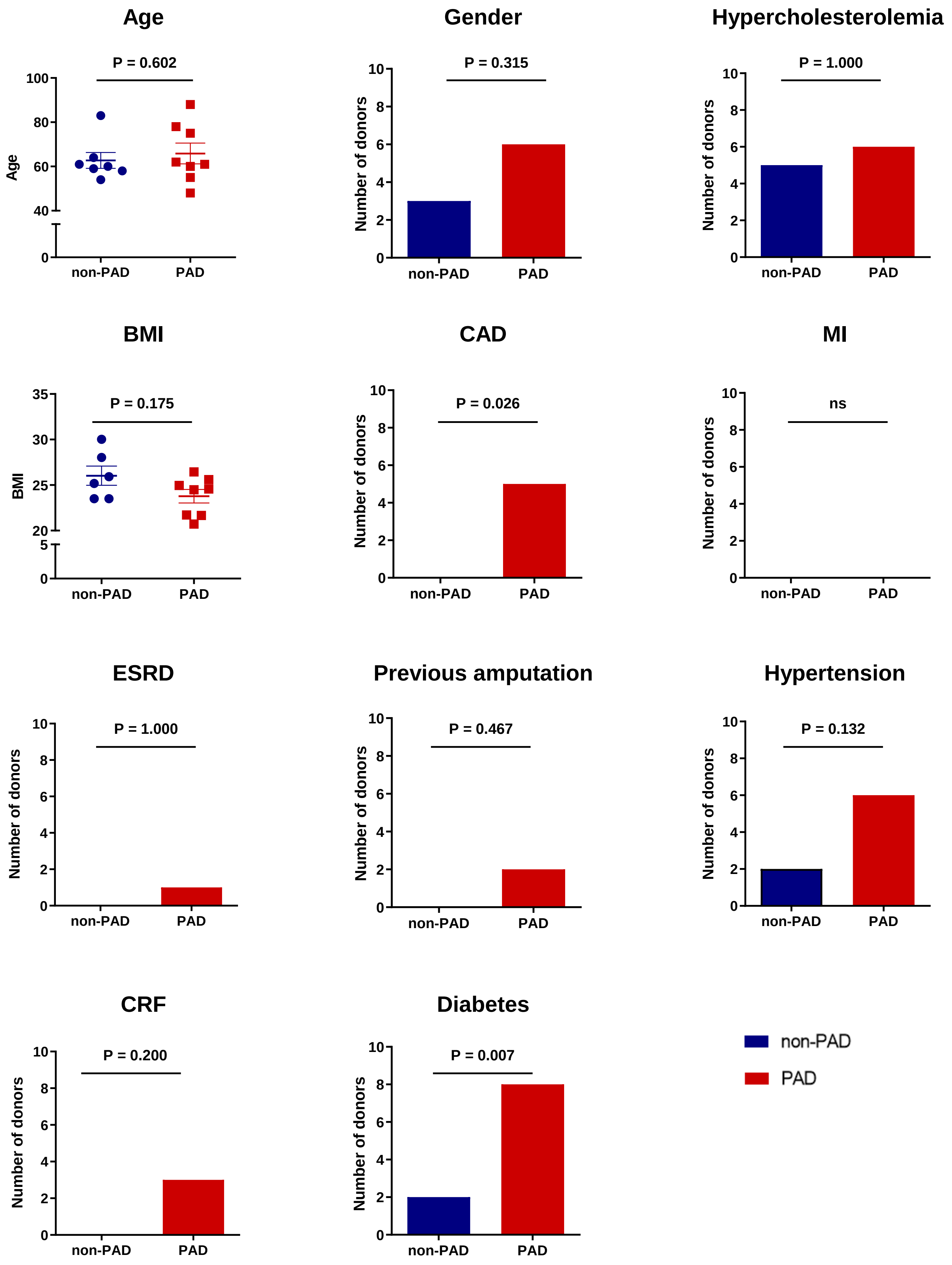

Age
P = 0.602
Age
100
80
60
40
0
non-PAD
PAD
Gender
P = 0.315
Number of donors
Hypercholesterolemia
P = 1.000
BMI
P = 0.175
35
30
25
20
5
CAD
P = 0.026
MI
ns
ESRD
P = 1.000
Previous amputation
P = 0.467
Hypertension
P = 0.132
CRF
P = 0.200
Diabetes
P = 0.007
non-PAD
PAD

**Figure S1. Demographic and Clinical Characteristics of Non-PAD Donors and PAD Patients**

Comparison of demographic and clinical characteristics between non-PAD and PAD donors. For age and body mass index (BMI), each symbol represents an individual donor, and horizontal lines and error bars indicate the mean ± SEM. Categorical variables are presented as the number of donors with each characteristic. Age and BMI were compared between groups using two-sided Mann–Whitney U tests, and categorical variables were compared using two-sided Fisher's exact tests. P values are shown for each comparison. n = 7 non-PAD donors and n = 8 PAD donors. Blue indicates non-PAD donors, and red indicates PAD donors. ns, not significant; PAD, peripheral artery disease; BMI, body mass index; SEM, standard error of the mean; CAD, coronary artery disease; MI, myocardial infarction; ESRD, end-stage renal disease; CRF, chronic renal failure.

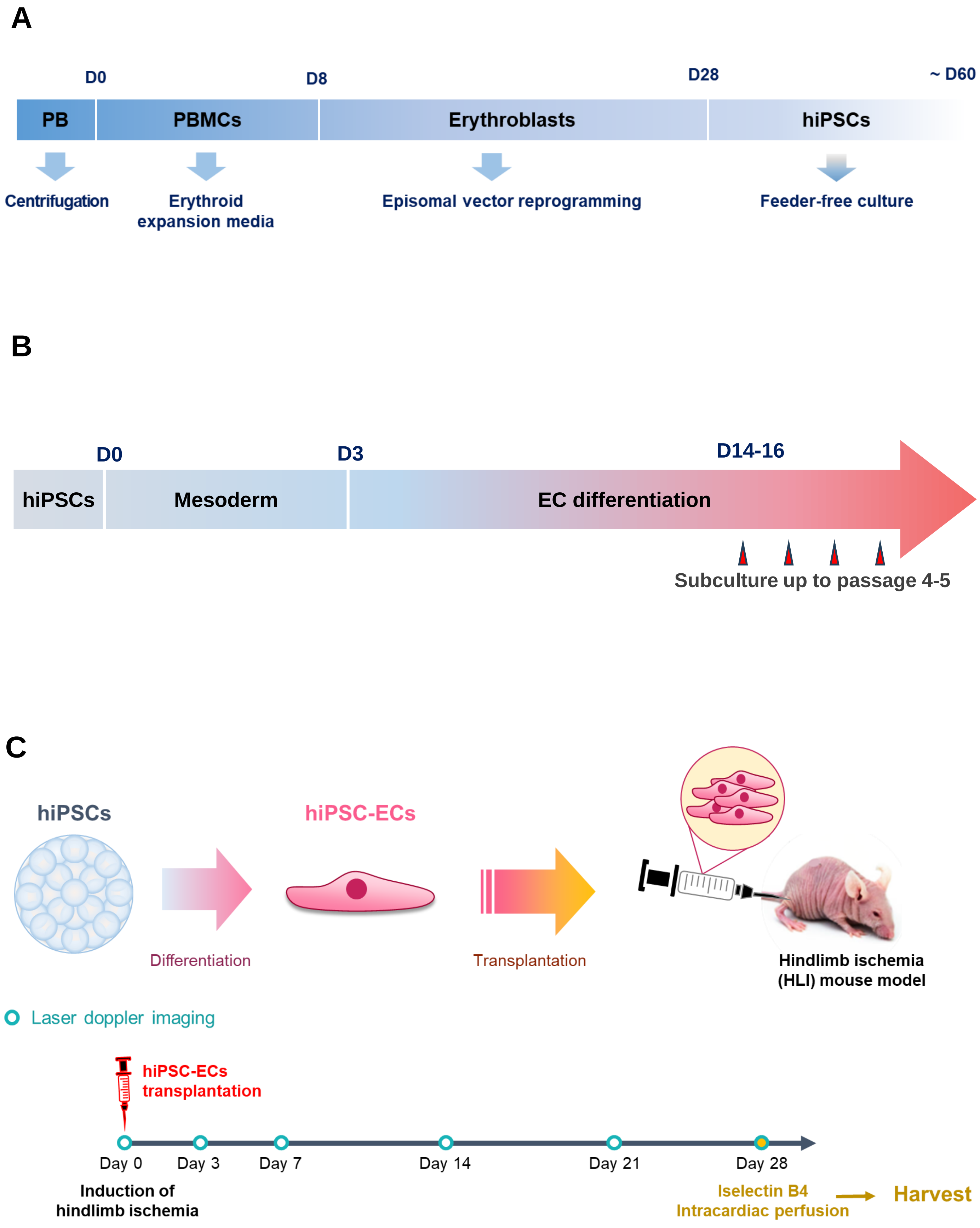
A
D0
D8
D28
~ D60
PB
PBMCs
Erythroblasts
hiPSCs
Centrifugation
Erythroid expansion media
Episomal vector reprogramming
Feeder-free culture
B
D0
D3
D14-16
hiPSCs
Mesoderm
EC differentiation
Subculture up to passage 4-5
C
hiPSCs
hiPSC-ECs
Differentiation
Transplantation
Hindlimb ischemia (HLI) mouse model
Laser doppler imaging
hiPSC-ECs transplantation
Day 0
Day 3
Day 7
Day 14
Day 21
Day 28
Induction of hindlimb ischemia
Iselectin B4 Intracardiac perfusion
Harvest

**Figure S2. Experimental Schematic of the Study**

**A**, Three stages of hiPSC generation: Stage 1, erythroblast enrichment; Stage 2, electrotransfection; and Stage 3, hiPSC initiation. **B**, Three stages of differentiation from hiPSCs into ECs: Stage 1, mesoderm induction; Stage 2, endothelial differentiation; and Stage 3, EC enrichment. **C**, Schematic overview of the in vivo experimental design.

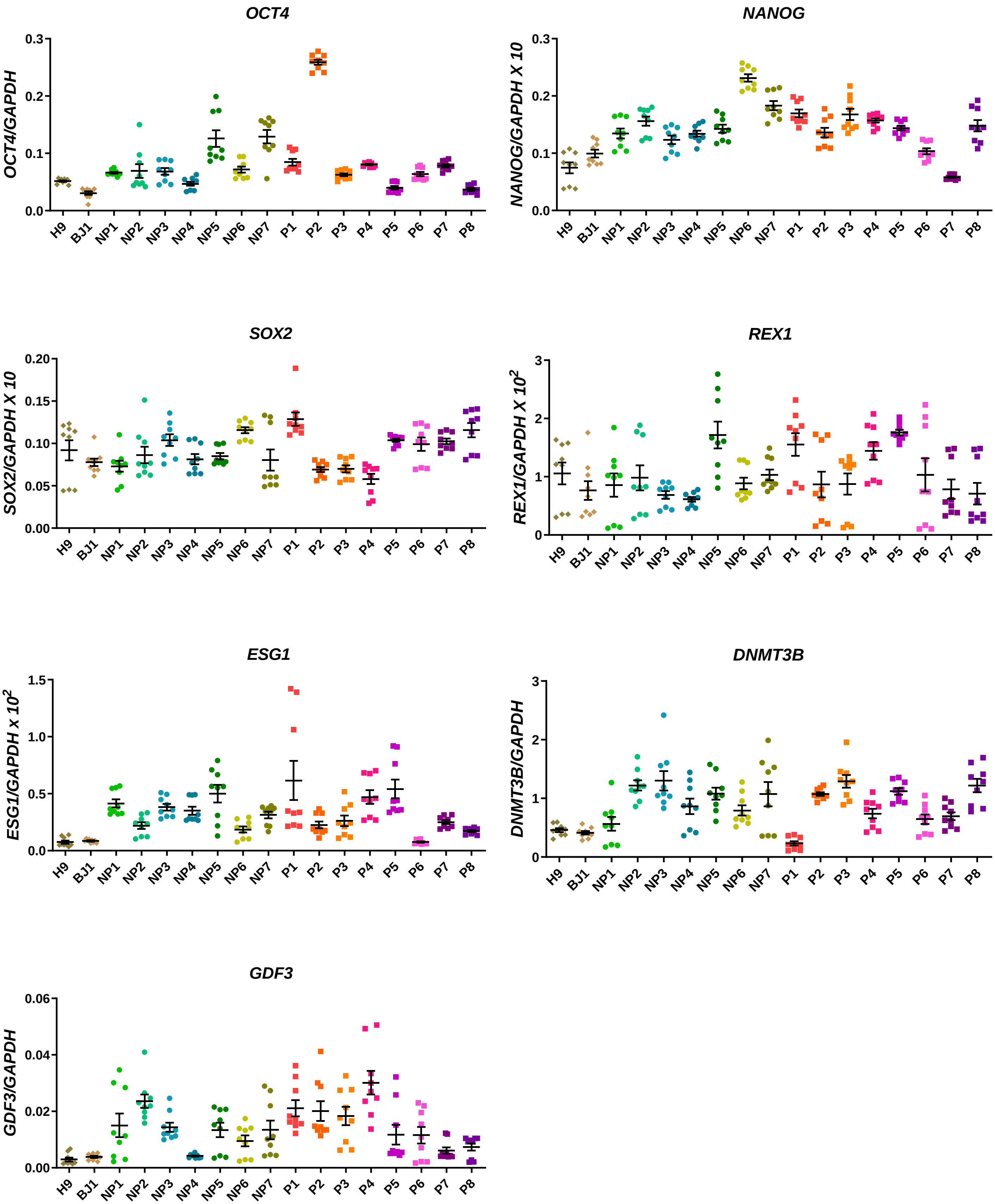

OCT4
OCT4/GAPDH
0.0
0.1
0.2
0.3
NANOG
NANOG/GAPDH X 10
0.0
0.1
0.2
0.3
SOX2
SOX2/GAPDH X 10
0.00
0.05
0.10
0.15
0.20
REX1
$REX1/GAPDH\ X\ 10^2$
0
1
2
3
ESG1
$ESG1/GAPDH\ x\ 10^2$
0.0
0.5
1.0
1.5
DNMT3B
DNMT3B/GAPDH
0
1
2
3
GDF3
GDF3/GAPDH
0.00
0.02
0.04
0.06
H9
BJ1
NP1
NP2
NP3
NP4
NP5
NP6
NP7
P1
P2
P3
P4
P5
P6
P7
P8

**Figure S3. mRNA Expression of Pluripotency-related Genes in Individual hiPSC Lines**

Relative mRNA expression in hiPSCs was normalized to GAPDH (set to 1). Data represent three independent experiments, each with technical triplicates per cell line (n = 7 for non-PAD; n = 8 for PAD). Data are shown as mean ± SEM.

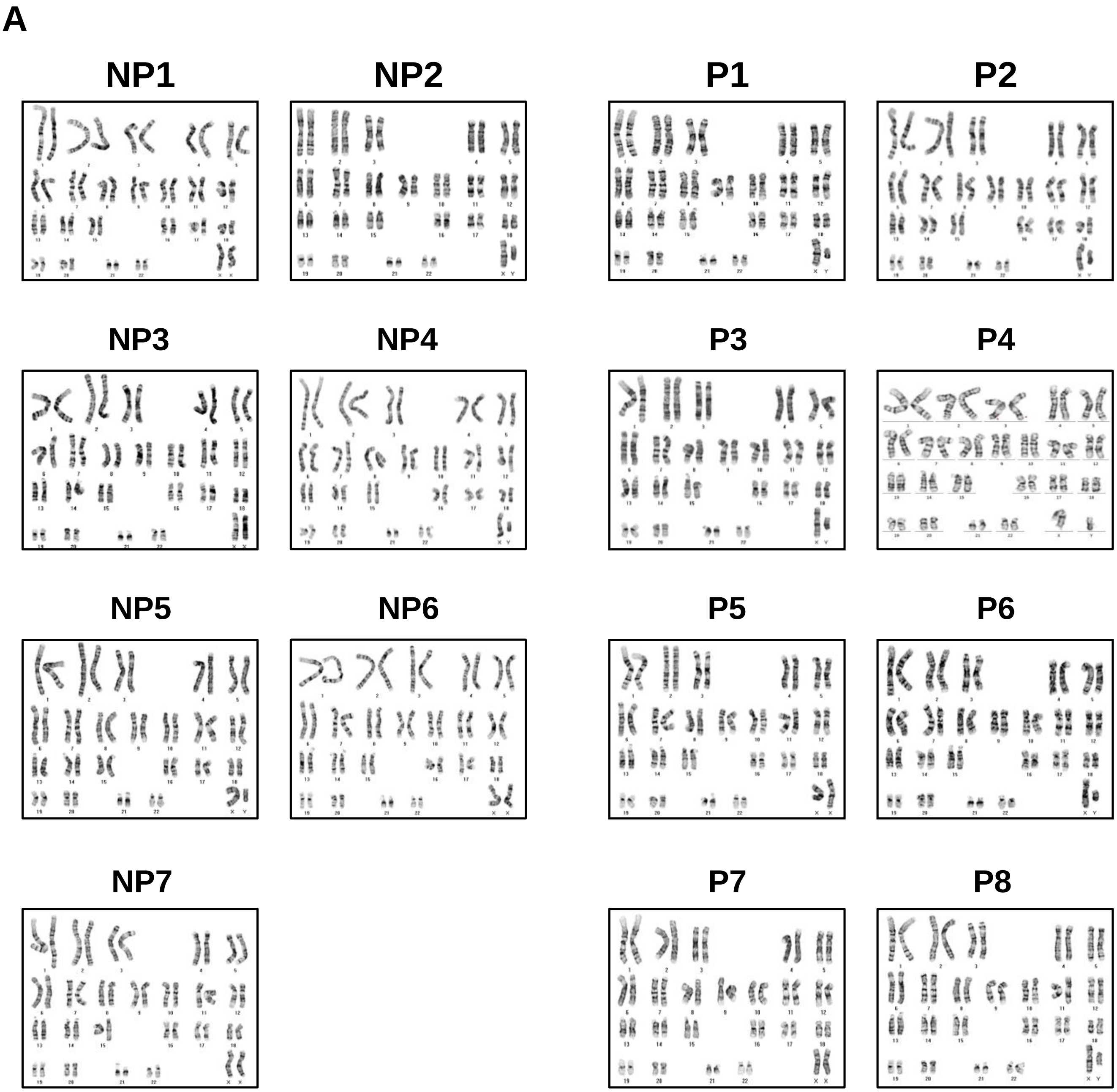
A
NP1
NP2
P1
P2
NP3
NP4
P3
P4
NP5
NP6
P5
P6
NP7
P7
P8

## B

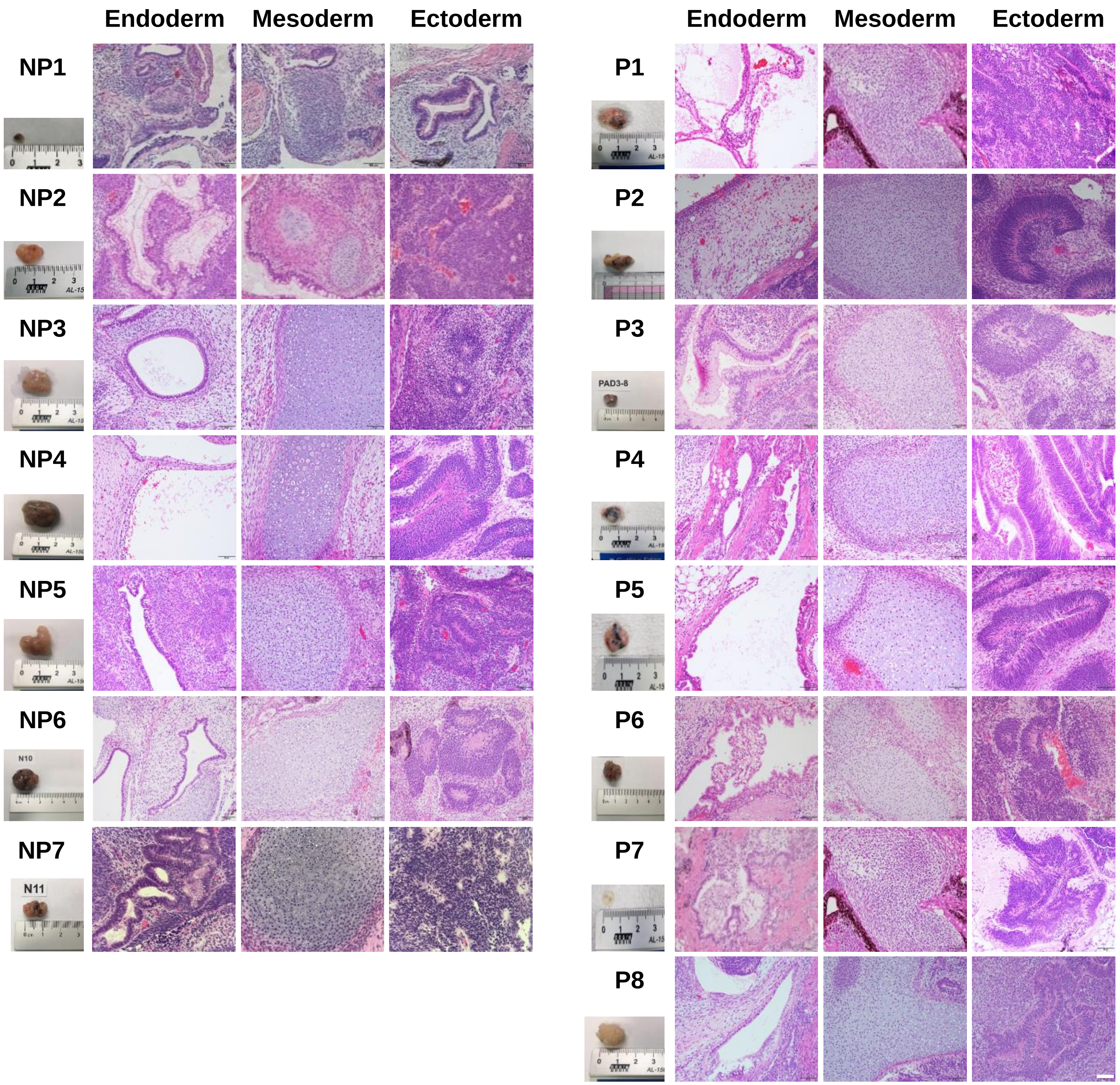
Endoderm
Mesoderm
Ectoderm
NP1
NP2
NP3
NP4
NP5
NP6
N10
NP7
N11
P1
P2
P3
PAD3-8
P4
P5
P6
P7
P8

**Figure S4. Characterization of hiPSCs Derived from non-PAD and PAD Donors**
**A**, Representative G-banding karyotype analysis of hiPSCs showing 23 pairs of human chromosomes. n = 7 for non-PAD; n = 8 for PAD. **B**, Teratoma formation assay in immunodeficient mice subcutaneously injected with $1 \times 10^7$ hiPSCs. Photomicrographs show representative teratomas collected 6–12 weeks post-injection, containing derivatives of all three germ layers: endoderm (gut epithelium, left), mesoderm (cartilage, middle), and ectoderm (neural tissue, right). For each hiPSC line, the three representative germ-layer images were obtained from the same teratoma. Tissue sections were stained with hematoxylin and eosin (H&E). Scale bars in B, 50 μm.

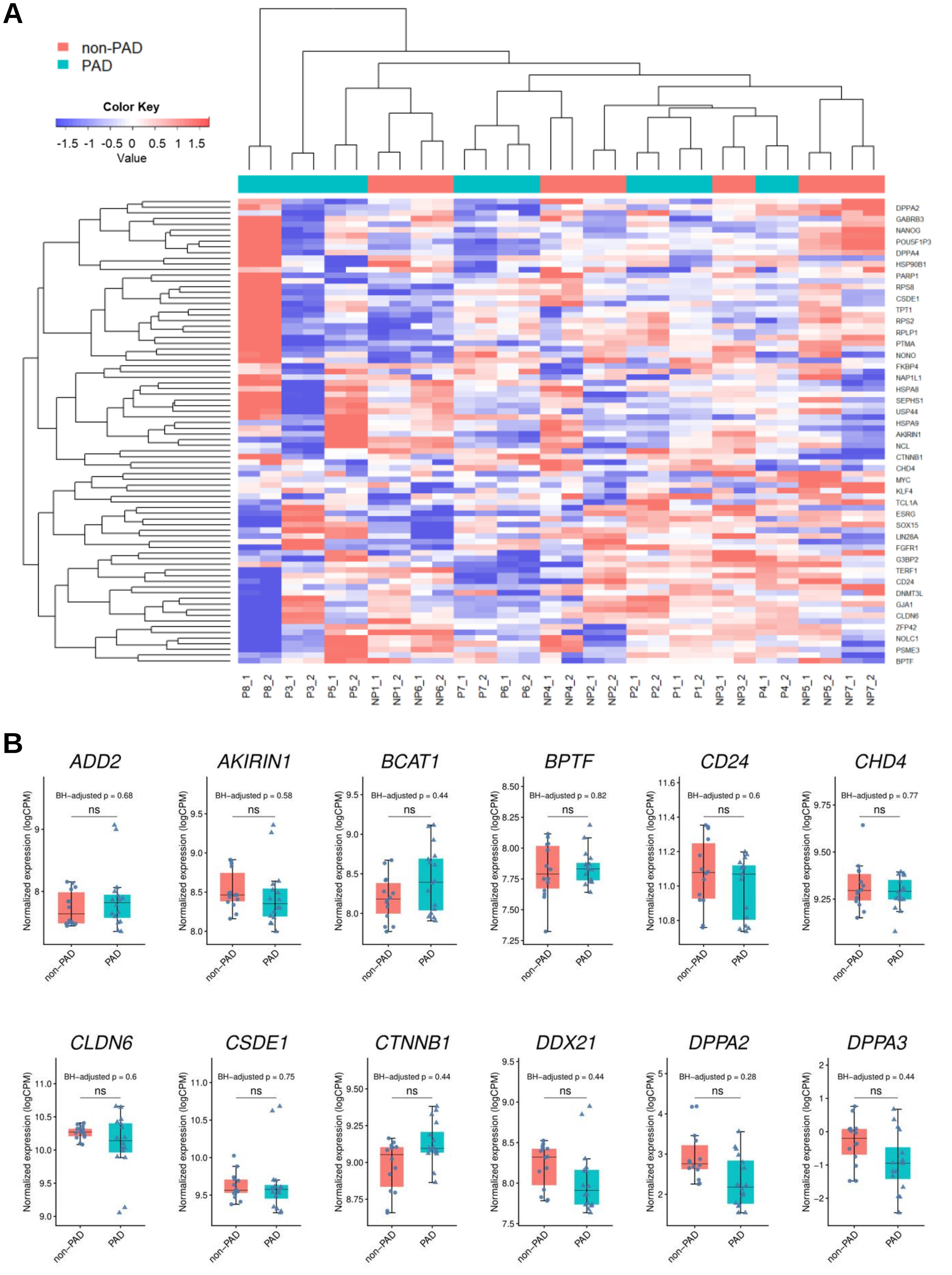
A
non-PAD
PAD
Color Key
-1.5 -1 -0.5 0 0.5 1 1.5
Value
DPPA2
GABRB3
NANOG
POU5F1P3
DPPA4
HSP90B1
PARP1
RPS8
CSDE1
TPT1
RPS2
RPLP1
PTMA
NONO
FKBP4
NAP1L1
HSPA8
SEPHS1
USP44
HSPA9
AKIRIN1
NCL
CTNNB1
CHD4
MYC
KLF4
TCL1A
ESRG
SOX15
LIN28A
FGFR1
G3BP2
TERF1
CD24
DNMT3L
GJA1
CLDN6
ZFP42
NOLC1
PSME3
BPTF
P8_1 P8_2 P3_1 P3_2 P5_1 P5_2 NP1_1 NP1_2 NP6_1 NP6_2 P7_1 P7_2 P6_1 P6_2 NP4_1 NP4_2 NP2_1 NP2_2 P2_1 P2_2 P1_1 P1_2 NP3_1 NP3_2 P4_1 P4_2 NP5_1 NP5_2 NP7_1 NP7_2
B
ADD2
BH-adjusted p = 0.68
AKIRIN1
BH-adjusted p = 0.58
BCAT1
BH-adjusted p = 0.44
BPTF
BH-adjusted p = 0.82
CD24
BH-adjusted p = 0.6
CHD4
BH-adjusted p = 0.77
CLDN6
BH-adjusted p = 0.6
CSDE1
BH-adjusted p = 0.75
CTNNB1
BH-adjusted p = 0.44
DDX21
BH-adjusted p = 0.44
DPPA2
BH-adjusted p = 0.28
DPPA3
BH-adjusted p = 0.44
ns
Normalized expression (logCPM)
non-PAD
PAD

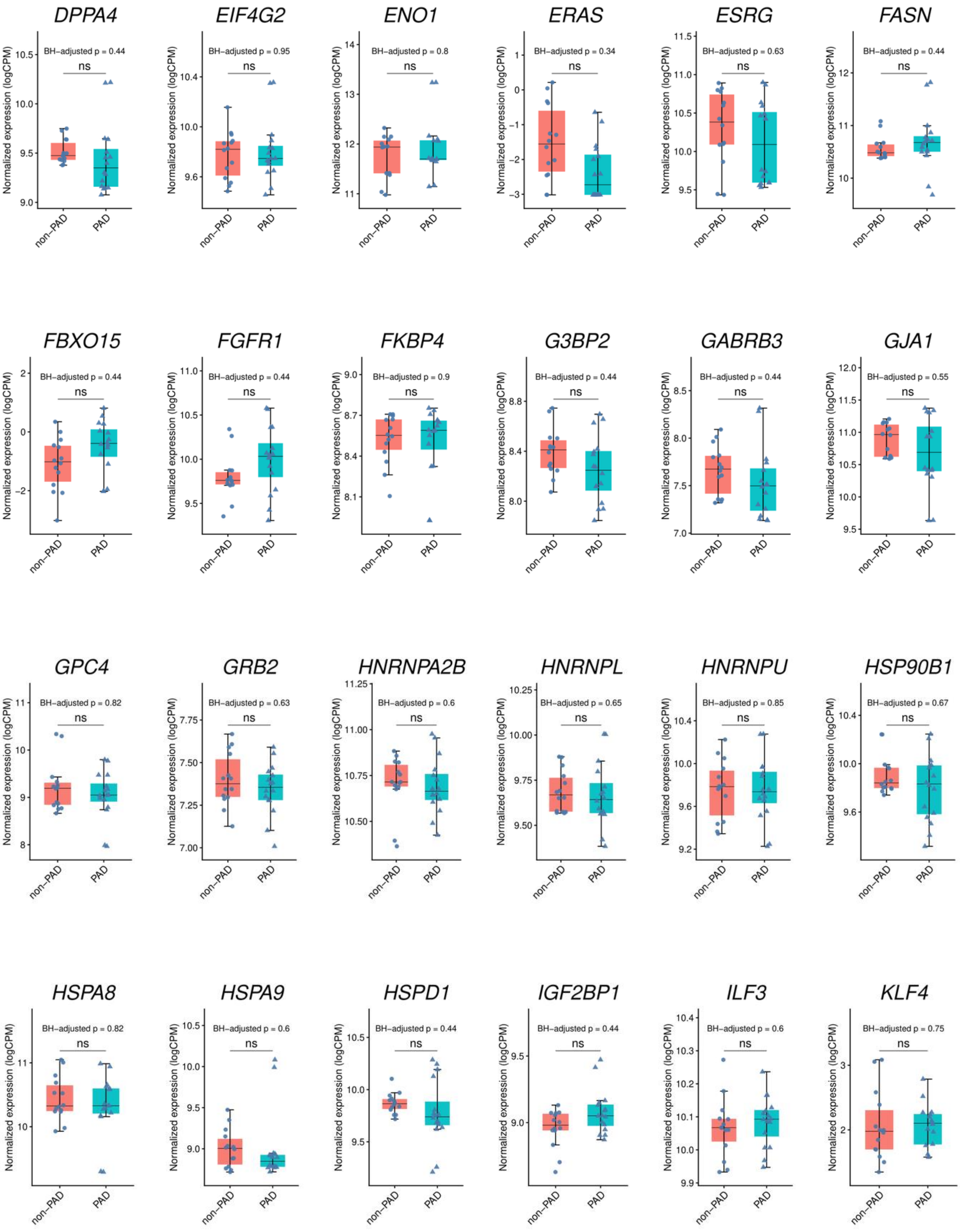
DPPA4
BH−adjusted p = 0.44
ns
EIF4G2
BH−adjusted p = 0.95
ns
ENO1
BH−adjusted p = 0.8
ns
ERAS
BH−adjusted p = 0.34
ns
ESRG
BH−adjusted p = 0.63
ns
FASN
BH−adjusted p = 0.44
ns
FBXO15
BH−adjusted p = 0.44
ns
FGFR1
BH−adjusted p = 0.44
ns
FKBP4
BH−adjusted p = 0.9
ns
G3BP2
BH−adjusted p = 0.44
ns
GABRB3
BH−adjusted p = 0.44
ns
GJA1
BH−adjusted p = 0.55
ns
GPC4
BH−adjusted p = 0.82
ns
GRB2
BH−adjusted p = 0.63
ns
HNRNPA2B
BH−adjusted p = 0.6
ns
HNRNPL
BH−adjusted p = 0.65
ns
HNRNPU
BH−adjusted p = 0.85
ns
HSP90B1
BH−adjusted p = 0.67
ns
HSPA8
BH−adjusted p = 0.82
ns
HSPA9
BH−adjusted p = 0.6
ns
HSPD1
BH−adjusted p = 0.44
ns
IGF2BP1
BH−adjusted p = 0.44
ns
ILF3
BH−adjusted p = 0.6
ns
KLF4
BH−adjusted p = 0.75
ns
Normalized expression (logCPM)
non−PAD
PAD

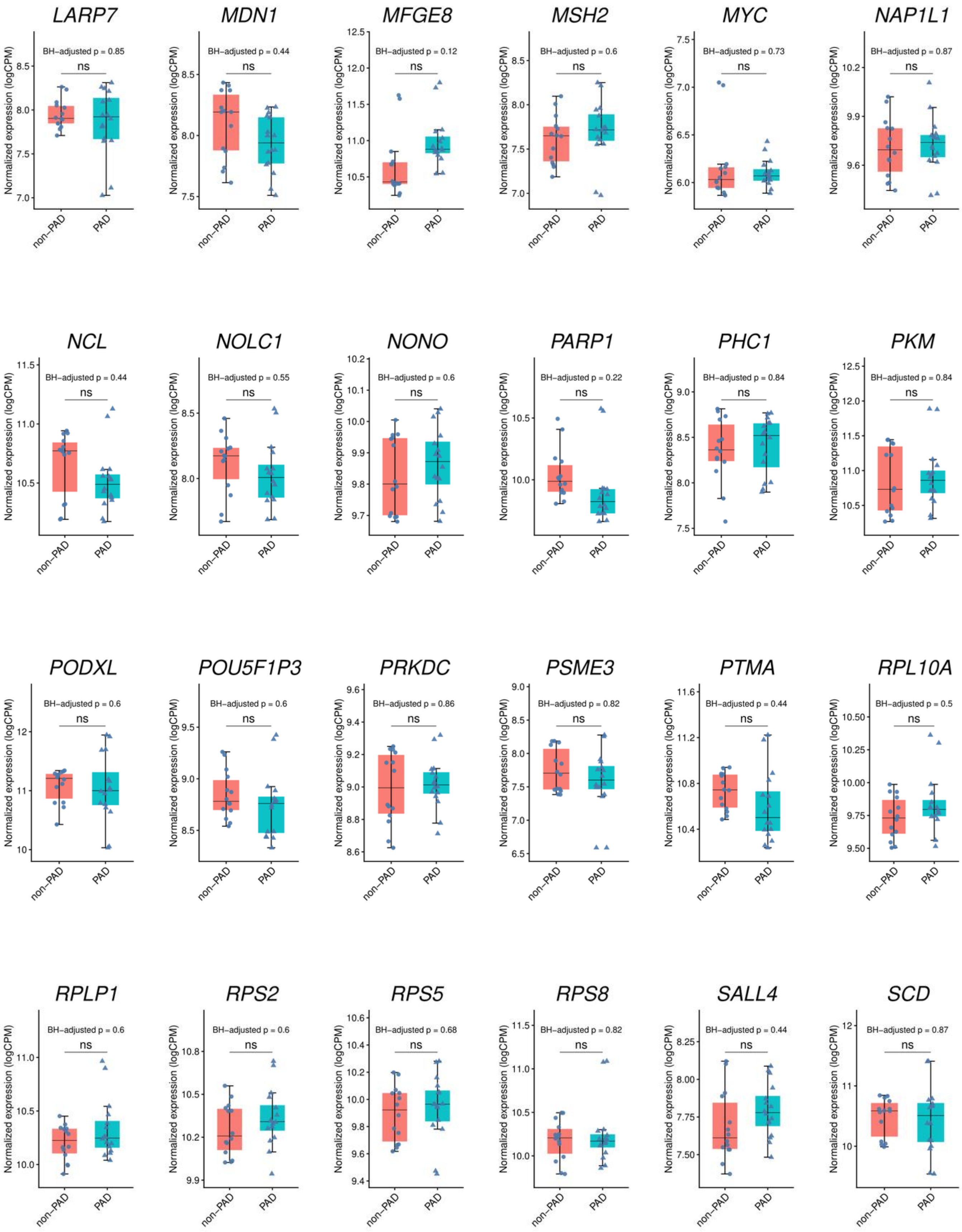
LARP7
BH-adjusted p = 0.85
MDN1
BH-adjusted p = 0.44
MFGE8
BH-adjusted p = 0.12
MSH2
BH-adjusted p = 0.6
MYC
BH-adjusted p = 0.73
NAP1L1
BH-adjusted p = 0.87
NCL
BH-adjusted p = 0.44
NOLC1
BH-adjusted p = 0.55
NONO
BH-adjusted p = 0.6
PARP1
BH-adjusted p = 0.22
PHC1
BH-adjusted p = 0.84
PKM
BH-adjusted p = 0.84
PODXL
BH-adjusted p = 0.6
POU5F1P3
BH-adjusted p = 0.6
PRKDC
BH-adjusted p = 0.86
PSME3
BH-adjusted p = 0.82
PTMA
BH-adjusted p = 0.44
RPL10A
BH-adjusted p = 0.5
RPLP1
BH-adjusted p = 0.6
RPS2
BH-adjusted p = 0.6
RPS5
BH-adjusted p = 0.68
RPS8
BH-adjusted p = 0.82
SALL4
BH-adjusted p = 0.44
SCD
BH-adjusted p = 0.87
ns
Normalized expression (logCPM)
non–PAD
PAD

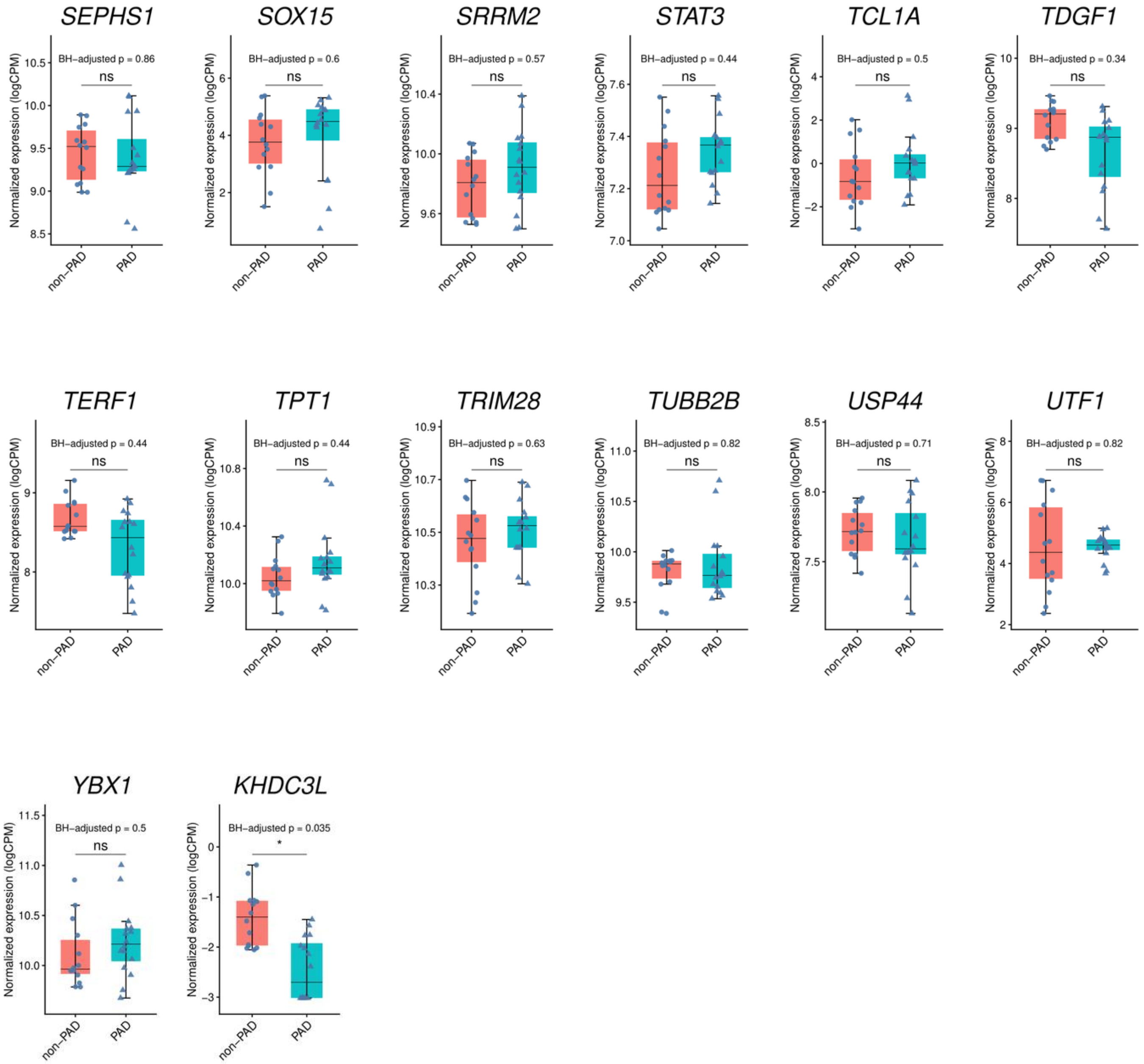
SEPHS1
BH−adjusted p = 0.86
ns
SOX15
BH−adjusted p = 0.6
ns
SRRM2
BH−adjusted p = 0.57
ns
STAT3
BH−adjusted p = 0.44
ns
TCL1A
BH−adjusted p = 0.5
ns
TDGF1
BH−adjusted p = 0.34
ns
TERF1
BH−adjusted p = 0.44
ns
TPT1
BH−adjusted p = 0.44
ns
TRIM28
BH−adjusted p = 0.63
ns
TUBB2B
BH−adjusted p = 0.82
ns
USP44
BH−adjusted p = 0.71
ns
UTF1
BH−adjusted p = 0.82
ns
YBX1
BH−adjusted p = 0.5
ns
KHDC3L
BH−adjusted p = 0.035
*
Normalized expression (logCPM)
non−PAD
PAD

**Figure S5. Expression Levels of Pluripotency-related Genes in hiPSCs**

**A**, Heatmap showing row-scaled, TMM-normalized $\log_2$ counts per million (logCPM) values for 82 pluripotency-related genes in hiPSCs derived from non-PAD (pink) and PAD (turquoise) donors. Red indicates relatively higher expression, and blue indicates relatively lower expression. **B**, Box plots showing TMM-normalized $\log_2$ counts per million (logCPM) values for the remaining 74 pluripotency-related genes, excluding the eight representative genes presented in Figure 2D. Genes are arranged alphabetically, with *KHDC3L* presented last. Each point represents one replicate sample (14 non-PAD samples from 7 donor-derived lines and 16 PAD samples from 8 donor-derived lines; two replicate samples per line). Boxes indicate the median and interquartile range, and whiskers extend to 1.5 times the interquartile range. P values were calculated using the two-sided Wilcoxon rank-sum test and adjusted for multiple comparisons using the Benjamini–Hochberg method. BH-adjusted p values are shown. ns, not significant; *BH-adjusted $p < 0.05$.

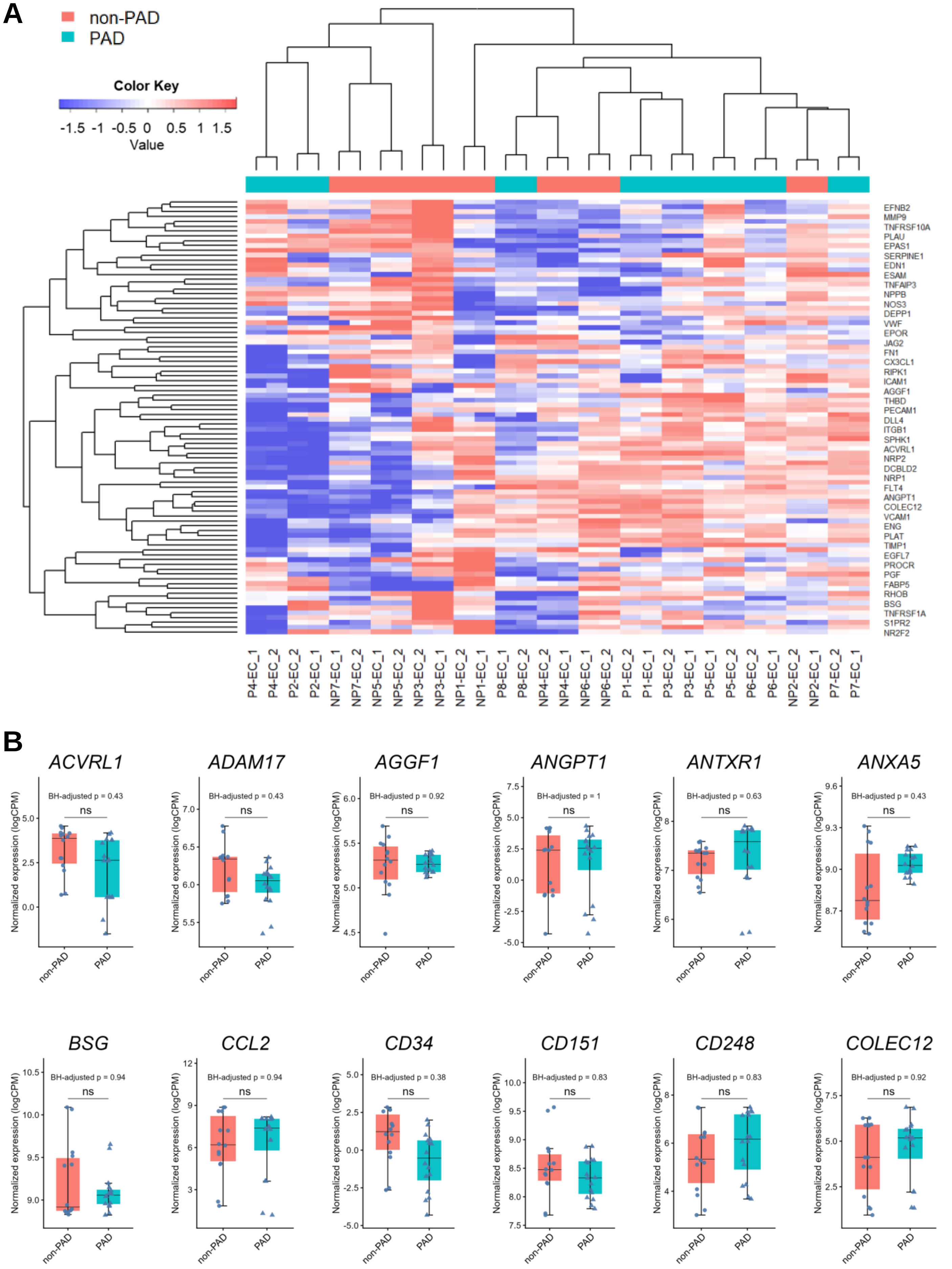
A
non-PAD
PAD
Color Key
-1.5 -1 -0.5 0 0.5 1 1.5
Value
EFNB2
MMP9
TNFRSF10A
PLAU
EPAS1
SERPINE1
EDN1
ESAM
TNFAIP3
NPPB
NOS3
DEPP1
VWF
EPOR
JAG2
FN1
CX3CL1
RIPK1
ICAM1
AGGF1
THBD
PECAM1
DLL4
ITGB1
SPHK1
ACVRL1
NRP2
DCBLD2
NRP1
FLT4
ANGPT1
COLEC12
VCAM1
ENG
PLAT
TIMP1
EGFL7
PROCR
PGF
FABP5
RHOB
BSG
TNFRSF1A
S1PR2
NR2F2
P4-EC_1
P4-EC_2
P2-EC_2
P2-EC_1
NP7-EC_1
NP7-EC_2
NP5-EC_1
NP5-EC_2
NP3-EC_2
NP3-EC_1
NP1-EC_2
NP1-EC_1
P8-EC_1
P8-EC_2
NP4-EC_2
NP4-EC_1
NP6-EC_1
NP6-EC_2
P1-EC_2
P1-EC_1
P3-EC_2
P3-EC_1
P5-EC_1
P5-EC_2
P6-EC_2
P6-EC_1
NP2-EC_2
NP2-EC_1
P7-EC_2
P7-EC_1
B
ACVRL1
BH-adjusted p = 0.43
ADAM17
BH-adjusted p = 0.43
AGGF1
BH-adjusted p = 0.92
ANGPT1
BH-adjusted p = 1
ANTXR1
BH-adjusted p = 0.63
ANXA5
BH-adjusted p = 0.43
BSG
BH-adjusted p = 0.94
CCL2
BH-adjusted p = 0.94
CD34
BH-adjusted p = 0.38
CD151
BH-adjusted p = 0.83
CD248
BH-adjusted p = 0.83
COLEC12
BH-adjusted p = 0.92
ns
Normalized expression (logCPM)
non-PAD
PAD

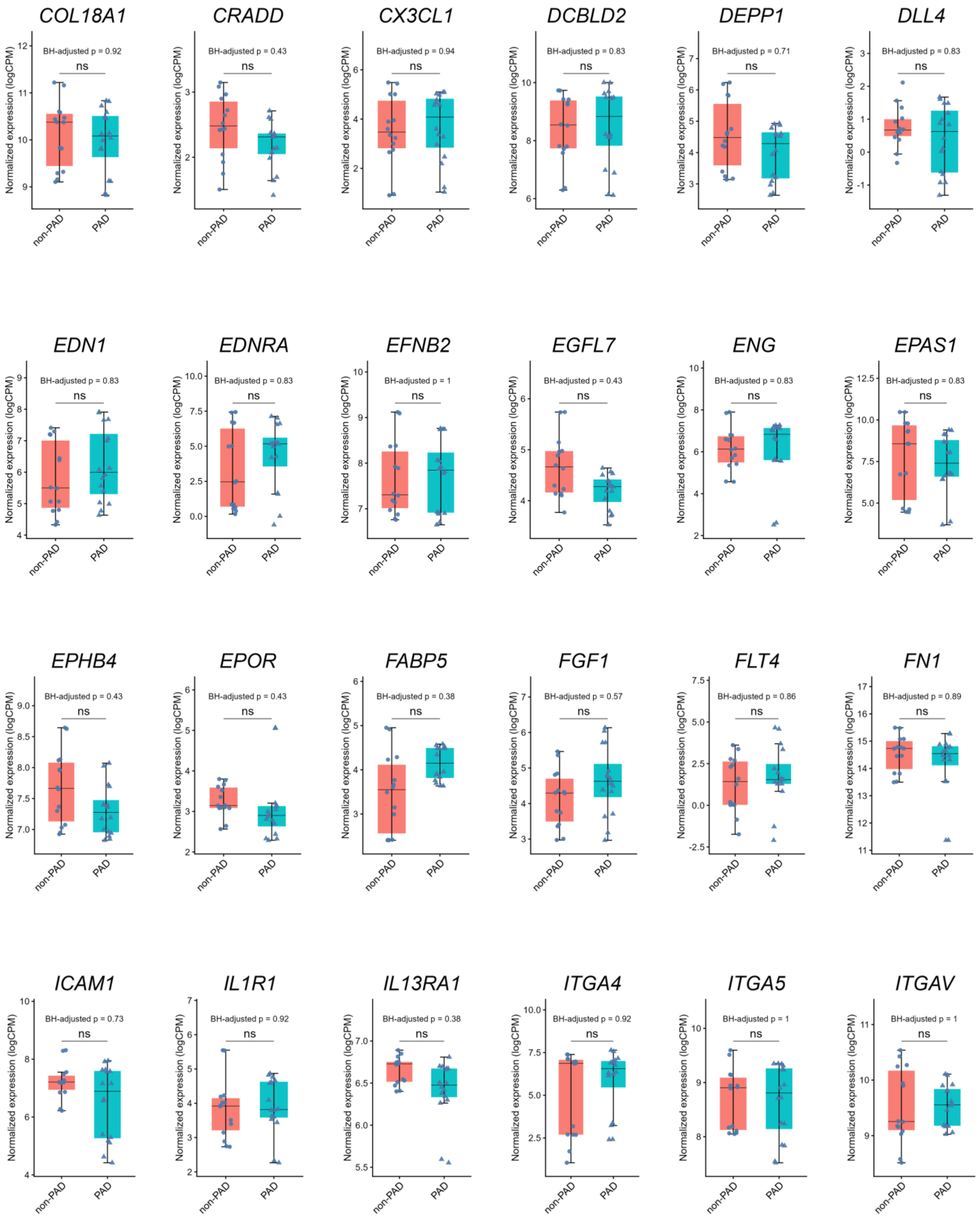
COL18A1
BH-adjusted p = 0.92
CRADD
BH-adjusted p = 0.43
CX3CL1
BH-adjusted p = 0.94
DCBLD2
BH-adjusted p = 0.83
DEPP1
BH-adjusted p = 0.71
DLL4
BH-adjusted p = 0.83
EDN1
BH-adjusted p = 0.83
EDNRA
BH-adjusted p = 0.83
EFNB2
BH-adjusted p = 1
EGFL7
BH-adjusted p = 0.43
ENG
BH-adjusted p = 0.83
EPAS1
BH-adjusted p = 0.83
EPHB4
BH-adjusted p = 0.43
EPOR
BH-adjusted p = 0.43
FABP5
BH-adjusted p = 0.38
FGF1
BH-adjusted p = 0.57
FLT4
BH-adjusted p = 0.86
FN1
BH-adjusted p = 0.89
ICAM1
BH-adjusted p = 0.73
IL1R1
BH-adjusted p = 0.92
IL13RA1
BH-adjusted p = 0.38
ITGA4
BH-adjusted p = 0.92
ITGA5
BH-adjusted p = 1
ITGAV
BH-adjusted p = 1
ns
Normalized expression (logCPM)
non-PAD
PAD

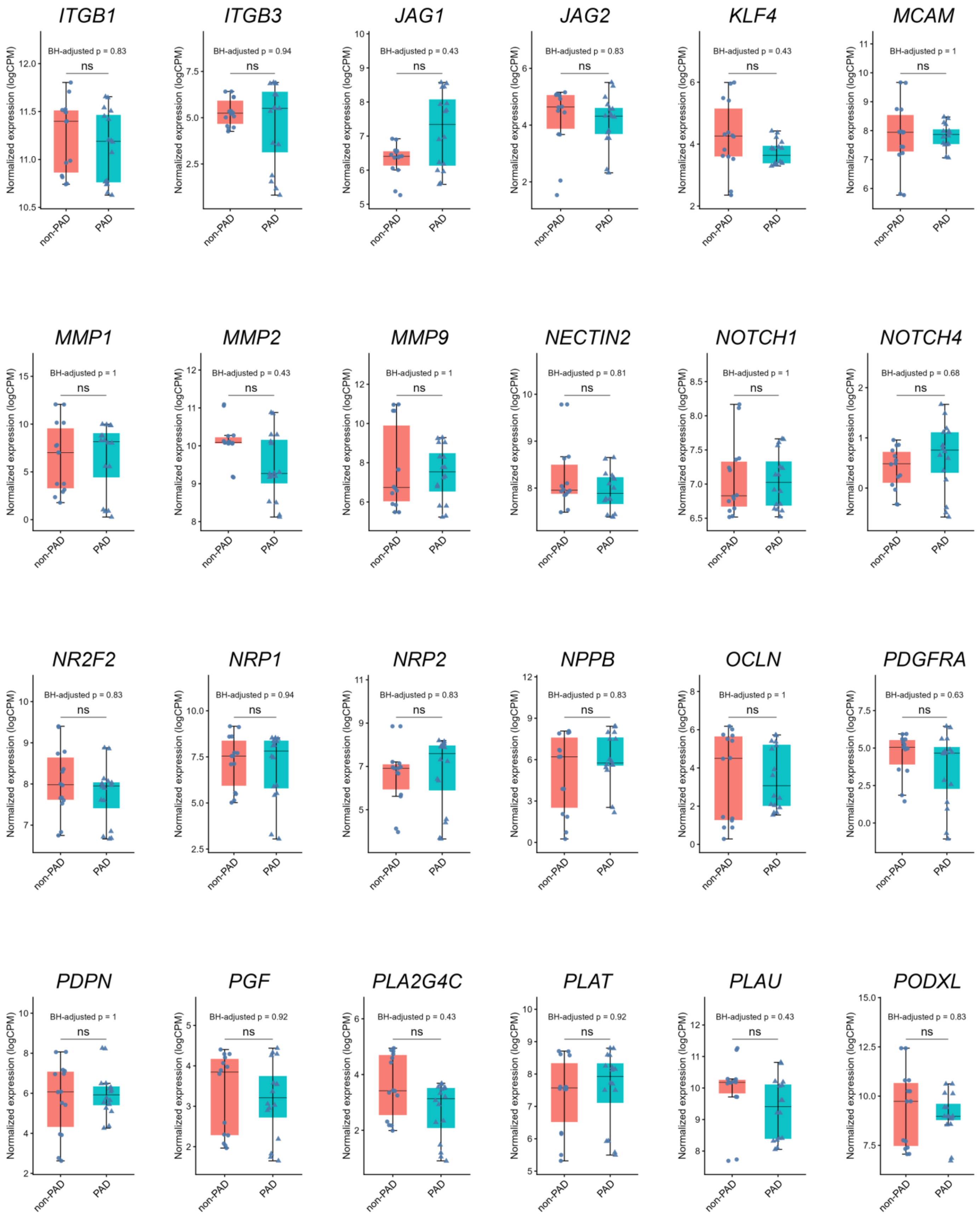
ITGB1
BH-adjusted p = 0.83
ITGB3
BH-adjusted p = 0.94
JAG1
BH-adjusted p = 0.43
JAG2
BH-adjusted p = 0.83
KLF4
BH-adjusted p = 0.43
MCAM
BH-adjusted p = 1
MMP1
BH-adjusted p = 1
MMP2
BH-adjusted p = 0.43
MMP9
BH-adjusted p = 1
NECTIN2
BH-adjusted p = 0.81
NOTCH1
BH-adjusted p = 1
NOTCH4
BH-adjusted p = 0.68
NR2F2
BH-adjusted p = 0.83
NRP1
BH-adjusted p = 0.94
NRP2
BH-adjusted p = 0.83
NPPB
BH-adjusted p = 0.83
OCLN
BH-adjusted p = 1
PDGFRA
BH-adjusted p = 0.63
PDPN
BH-adjusted p = 1
PGF
BH-adjusted p = 0.92
PLA2G4C
BH-adjusted p = 0.43
PLAT
BH-adjusted p = 0.92
PLAU
BH-adjusted p = 0.43
PODXL
BH-adjusted p = 0.83
ns
Normalized expression (logCPM)
non-PAD
PAD

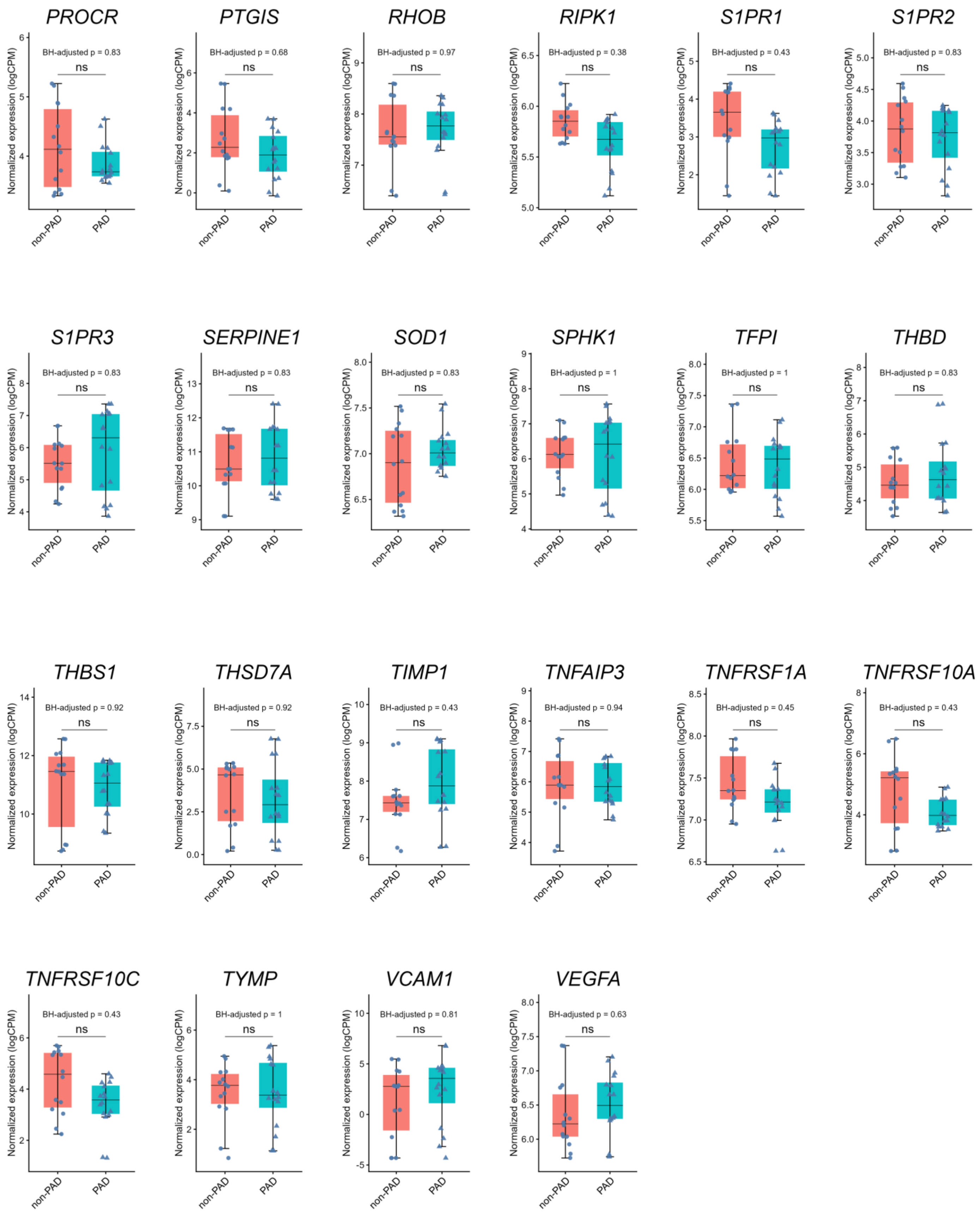
PROCR
BH-adjusted p = 0.83
PTGIS
BH-adjusted p = 0.68
RHOB
BH-adjusted p = 0.97
RIPK1
BH-adjusted p = 0.38
S1PR1
BH-adjusted p = 0.43
S1PR2
BH-adjusted p = 0.83
S1PR3
BH-adjusted p = 0.83
SERPINE1
BH-adjusted p = 0.83
SOD1
BH-adjusted p = 0.83
SPHK1
BH-adjusted p = 1
TFPI
BH-adjusted p = 1
THBD
BH-adjusted p = 0.83
THBS1
BH-adjusted p = 0.92
THSD7A
BH-adjusted p = 0.92
TIMP1
BH-adjusted p = 0.43
TNFAIP3
BH-adjusted p = 0.94
TNFRSF1A
BH-adjusted p = 0.45
TNFRSF10A
BH-adjusted p = 0.43
TNFRSF10C
BH-adjusted p = 0.43
TYMP
BH-adjusted p = 1
VCAM1
BH-adjusted p = 0.81
VEGFA
BH-adjusted p = 0.63
ns
Normalized expression (logCPM)
non-PAD
PAD

**Figure S6. Expression Levels of EC–related Genes in hiPSC-ECs**

**A**, Heatmap showing row-scaled, TMM-normalized $log_2$ counts per million (logCPM) values for 90 endothelial cell–related genes in hiPSC-ECs derived from non-PAD (pink) and PAD (turquoise) donors. Red indicates relatively higher expression, and blue indicates relatively lower expression. **B**, Box plots showing TMM-normalized $log_2$ counts per million (logCPM) values for the remaining 82 endothelial cell–related genes, excluding the eight representative genes presented in Figure 4D. Genes are arranged alphabetically. Each point represents one replicate sample (14 non-PAD samples from 7 donor-derived lines and 16 PAD samples from 8 donor-derived lines; two replicate samples per line). Boxes indicate the median and interquartile range, and whiskers extend to 1.5 times the interquartile range. P values were calculated using the two-sided Wilcoxon rank-sum test and adjusted for multiple comparisons using the Benjamini–Hochberg method. BH-adjusted p values are shown. ns, not significant.

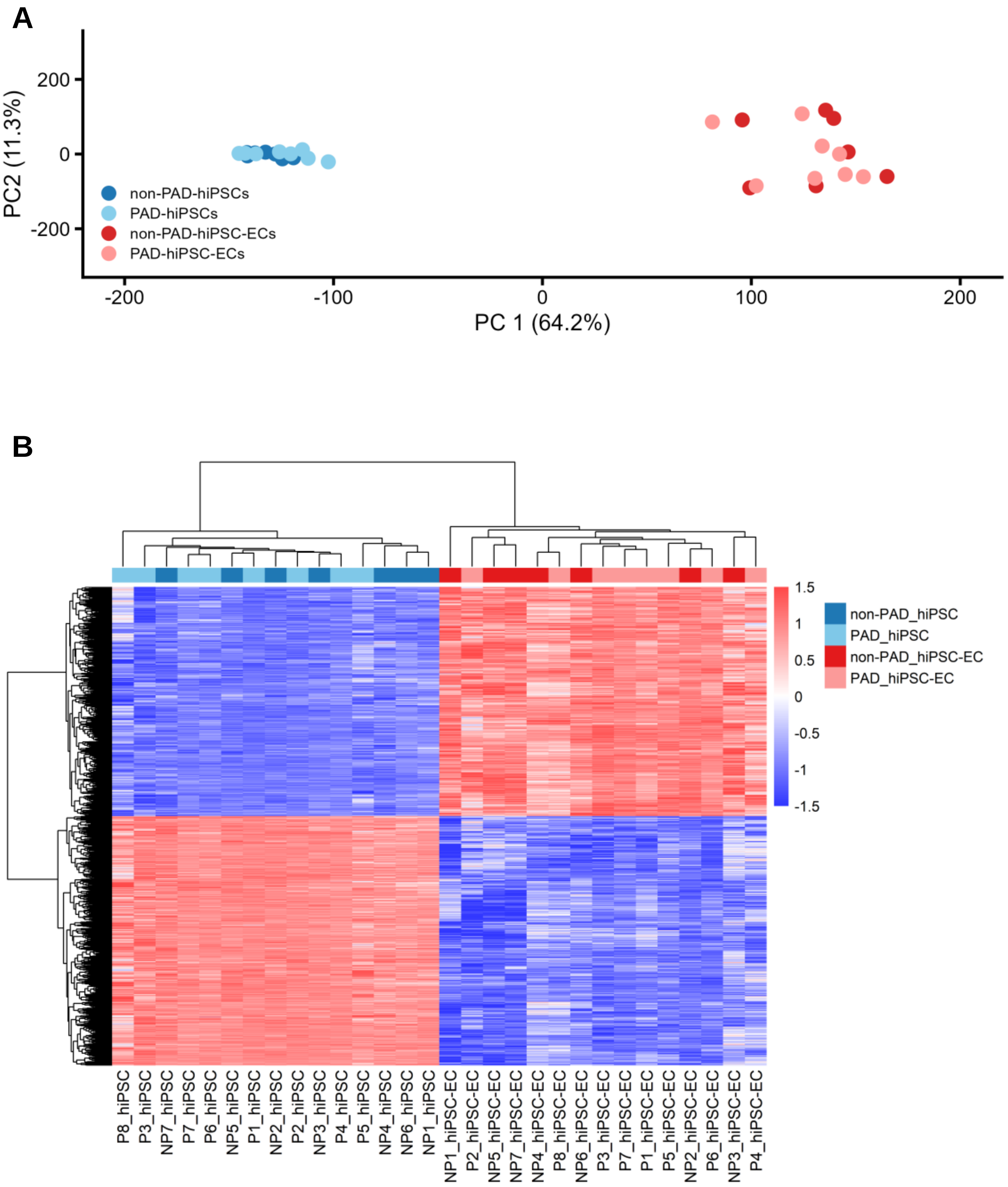
A
PC2 (11.3%)
200
0
-200
PC 1 (64.2%)
-200
-100
0
100
200
non-PAD-hiPSCs
PAD-hiPSCs
non-PAD-hiPSC-ECs
PAD-hiPSC-ECs
B
1.5
1
0.5
0
-0.5
-1
-1.5
non-PAD_hiPSC
PAD_hiPSC
non-PAD_hiPSC-EC
PAD_hiPSC-EC
P8_hiPSC
P3_hiPSC
NP7_hiPSC
P7_hiPSC
P6_hiPSC
NP5_hiPSC
P1_hiPSC
NP2_hiPSC
P2_hiPSC
NP3_hiPSC
P4_hiPSC
P5_hiPSC
NP4_hiPSC
NP6_hiPSC
NP1_hiPSC
NP1_hiPSC-EC
P2_hiPSC-EC
NP5_hiPSC-EC
NP7_hiPSC-EC
NP4_hiPSC-EC
P8_hiPSC-EC
NP6_hiPSC-EC
P3_hiPSC-EC
P7_hiPSC-EC
P1_hiPSC-EC
P5_hiPSC-EC
NP2_hiPSC-EC
P6_hiPSC-EC
NP3_hiPSC-EC
P4_hiPSC-EC

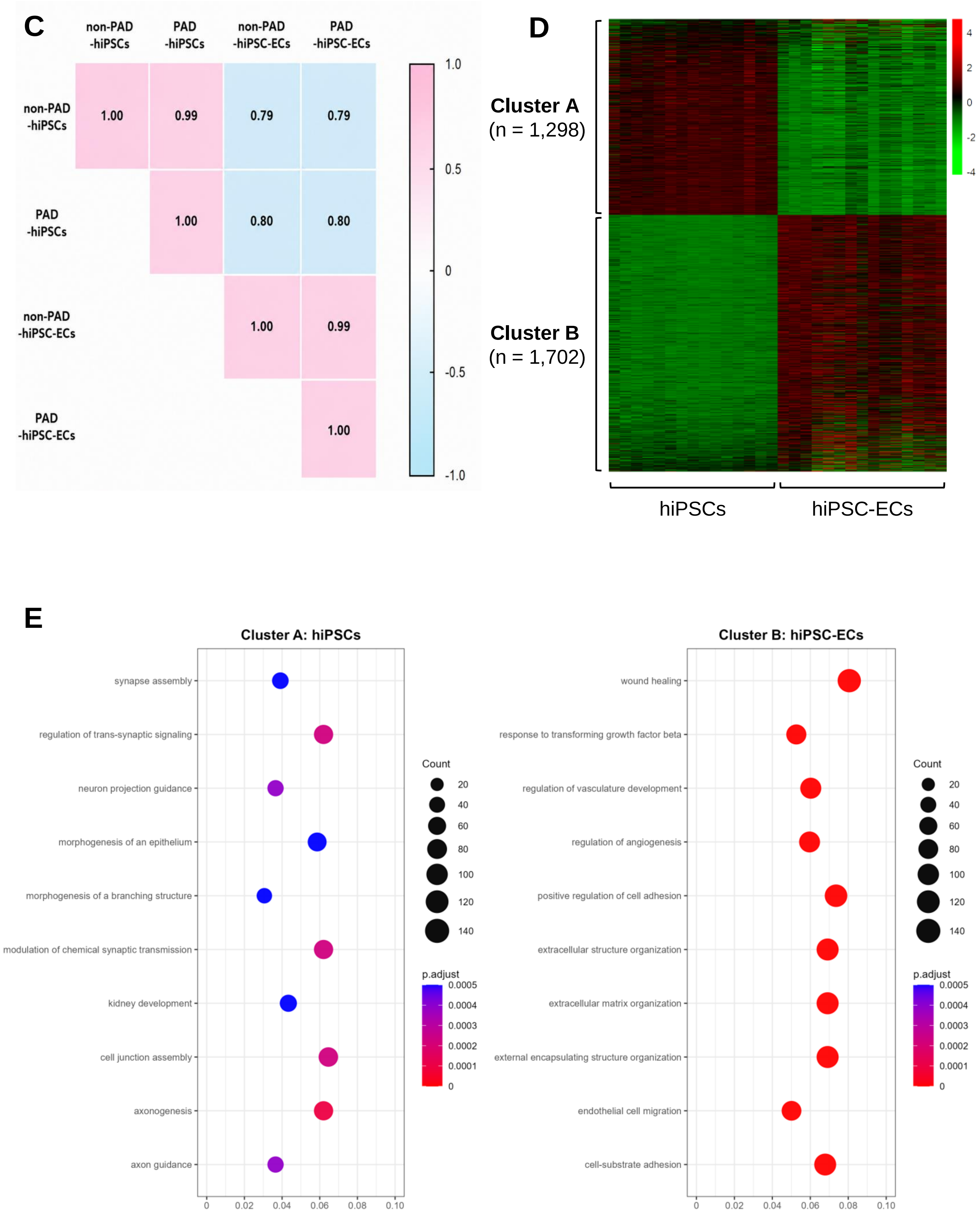
C
non-PAD -hiPSCs
PAD -hiPSCs
non-PAD -hiPSC-ECs
PAD -hiPSC-ECs
1.00
0.99
0.79
0.79
1.00
0.80
0.80
1.00
0.99
1.00
1.0
0.5
0
-0.5
-1.0
D
Cluster A
(n = 1,298)
Cluster B
(n = 1,702)
hiPSCs
hiPSC-ECs
4
2
0
-2
-4
E
Cluster A: hiPSCs
synapse assembly
regulation of trans-synaptic signaling
neuron projection guidance
morphogenesis of an epithelium
morphogenesis of a branching structure
modulation of chemical synaptic transmission
kidney development
cell junction assembly
axonogenesis
axon guidance
0
0.02
0.04
0.06
0.08
0.10
Count
20
40
60
80
100
120
140
p.adjust
0.0005
0.0004
0.0003
0.0002
0.0001
0
Cluster B: hiPSC-ECs
wound healing
response to transforming growth factor beta
regulation of vasculature development
regulation of angiogenesis
positive regulation of cell adhesion
extracellular structure organization
extracellular matrix organization
external encapsulating structure organization
endothelial cell migration
cell-substrate adhesion
0
0.02
0.04
0.06
0.08
0.10
Count
20
40
60
80
100
120
140
p.adjust
0.0005
0.0004
0.0003
0.0002
0.0001
0

**Figure S7. Comparative transcriptomic analyses of hiPSCs and hiPSC-ECs by RNA-seq.**

**A**, PCA of hiPSCs derived from non-PAD (blue) and PAD (sky blue) donors, and hiPSC-ECs derived from non-PAD (red) and PAD (pink) donors, demonstrating clear separation according to cell type. **B**, Hierarchical clustering heatmap of hiPSCs and hiPSC-ECs derived from non-PAD and PAD donors (n = 7 for non-PAD and n = 8 for PAD in both hiPSCs and hiPSC-ECs). Samples clustered primarily according to cell type rather than PAD status. Red indicates relatively higher expression, and blue indicates relatively lower expression. **C**, Pearson correlation matrix analysis demonstrating strong correlations within the same cell type irrespective of PAD status. **D**, K-means clustering analysis of the top 3,000 differentially expressed genes between hiPSCs and hiPSC-ECs, identifying distinct gene clusters associated with pluripotency and endothelial differentiation. **E**, Gene Ontology (GO) Biological Process enrichment analysis of genes enriched in each cluster identified by K-means clustering.

**Table S1.** Demographic Characteristics of Non-PAD Donors and PAD Patients Used for hiPSC Generation

Summary of demographic and clinical information for non-PAD donors and PAD patients from whom peripheral blood samples were collected for the generation of hiPSC lines.

| | non-PAD | PAD | P value |
|---|---|---|---|
| Age | 62.7 | 65.9 | 0.602 |
| Gender (Male) | 3 | 6 | 0.315 |
| Hypercholesterolemia | 5 | 6 | 1.000 |
| BMI | 26.0 | 23.8 | 0.175 |
| CAD | 0 | 5 | 0.026 |
| MI | 0 | 0 | NA |
| ESRD | 0 | 1 | 1.000 |
| Previous amputation | 0 | 2 | 0.467 |
| Hypertension | 2 | 6 | 0.132 |
| CRF | 0 | 3 | 0.200 |
| Diabetes | 2 | 8 | 0.007 |
| Donor | 7 | 8 | - |

**Table S2.** Primer Sequences Used in This Study
List of primer sequences used for quantitative RT–PCR analysis of pluripotency- and endothelial-related genes.

| | Forward | Reverse |
|---|---|---|
| *OCT4* | GTTTGTGCCAGGGTTTTTGGG | CAACCAGTTGCCCCAAACTCC |
| *NANOG* | GACTGTCTCTCCTCTTCCTTCCTC | TGCGACACTCTTCTCTGCAG |
| *SOX2* | CAGCAAATGACAGCTGCAAAAGAG | ATCTCTCTCATAAAAGTTTTCTTGTCGGC |
| *DNMT3B* | TGCTGCTCACAGGGCCCGATACTTC | TCCTTTCGAGCTCAGTGCACCACAAAAC |
| *REX1* | CAGATCCTAAACAGCTCGCAGAAT | GCGTACGCAAATTAAAGTCCAGA |
| *GDF3* | CTTATGCTACGTAAAGGAGCTGGG | GTGCCAACCCAGGTCCCGGAAGTT |
| *ESG1* | ATATCCCGCCGTGGGTGAAAGTTC | ACTCAGCCATGGACTGGAGCATCC |
| *CDH5* | CTGGCCATGGACCCTGATG | CGGAAGAACTGGCCCTTGT |
| *PECAM1* | TGTATTTCAAGACCTCTGTGCACTT | TTAGCCTGAGGAATTGCTGTGTT |
| *KDR* | ATCTCAATGTGGTCAACCTTCTAGGT | AAATTTGCAGAATTCCACAATCAC |
| *TEK* | CCTGCCTGACTGTGCTGTTG | TGCACATTTGCCCTCTTCAA |
| *VWF* | TGAGCCCCACCACTCTGTATG | GTAGCCTGCTGCAGTAGAAATCG |
| *NOS3* | CGGCATCACCAGGAAGAAGA | CATGAGCGAGGCGGAGAT |
| *OCT4* | GAACCGAGTGAGAGGCAACCT | TCTGCTGCAGTGTGGGTTTC |
| *oriP* | TTCCACGAGGGTAGTGAACC | TCGGGGGTGTTAGAGACAAC |
| *EBNA-1* | ATCGTCAAAGCTGCACACAG | CCCAGGAGTCCCAGTAGTCA |
| *GAPDH* | CAGTCCATGCCATCACTGCC | GGATGATGTTCTGGAGAGCCCC |

**Table S3**. List of Pluripotency-related Genes Analyzed in This Study
Comprehensive list of pluripotency-related genes analyzed in RNA-seq.

| Symbol | Name | Alias |
|---|---|---|
| ADD2 | adducin 2 | ADDB |
| AKIRIN1 | akirin 1 | 6330407G11Rik |
| BCAT1 | branched chain amino acid transaminase 1 | BCATC, BCT1, ECA39, MECA39, PNAS121, PP18 |
| BPTF | bromodomain PHD finger transcription factor | FAC1, FALZ, NEDDFL, NURF301 |
| CD24 | CD24 molecule | CD24A |
| CHD4 | chromodomain helicase DNA binding protein 4 | CHD-4, Mi-2b, Mi2-BETA, SIHIWES |
| CLDN6 | claudin 6 | |
| CSDE1 | cold shock domain containing E1 | D1S155E, UNR |
| CTNNB1 | catenin beta 1 | VR7, CTNNB, MRD19, NEDSDV, armadillo |
| DDX21 | DExD-box helicase 21 | GUA, GURDB, II/Gu, RH, RH II/Gu, RH-II/GU, RH-II/GuA, gu-alpha |
| DNMT3B | DNA methyltransferase 3 beta | ICF, ICF1, FSHD4, M.HsaIIIB |
| DPPA2 | developmental pluripotency associated 2 | CT100, ECAT15-2, PESCRG1 |
| DPPA3 | developmental pluripotency associated 3 | Pgc7, STELLA |
| DPPA4 | developmental pluripotency associated 4 | 2410091M23Rik |
| EIF4G2 | eukaryotic translation initiation factor 4 gamma 2 | AAG1, DAP5, NAT1, P97 |
| ENO1 | enolase 1 | ENO1L1, HEL-S-17, MPB1, NNE, PPH |
| ERAS | ES cell expressed Ras | HRAS2, HRASP |
| ESG1 | embryonal stem cell-specific gene 1 | DPPA5 |
| ESRG | embryonic stem cell related | HESRG |
| FASN | fatty acid synthase | FAS, OA-519, SDR27X1 |
| FBXO15 | F-box protein 15 | FBX15 |
| FGFR1 | fibroblast growth factor receptor 1 | BFGFR, CD331, CEK, ECCL, FGFBR |
| FKBP4 | FKBP prolyl isomerase 4 | FKBP51, FKBP52, FKBP59, HBI, Hsp56, PPIase, p52 |
| G3BP2 | G3BP stress granule assembly factor 2 | |
| GABRB3 | gamma-aminobutyric acid type A receptor subunit beta3 | DEE43, ECA5, EIEE43 |
| GDF3 | growth differentiation factor 3 | KFS3, MCOP7, MCOPCB6 |
| GJA1 | gap junction protein alpha 1 | AVSD3, CMDR, CX43, EKVP, EKVP3 |
| GPC4 | glypican 4 | K-glypican, KPTS |
| GRB2 | growth factor receptor bound protein 2 | ASH, EGFRBP-GRB2, Grb3-3, MST084, MSTP084, NCKAP2 |
| HNRNPA2B1 | heterogeneous nuclear ribonucleoprotein A2/B1 | HNRNPA2, HNRNPB1, HNRPA2, HNRPA2B1 |

**Table S3**. List of Pluripotency-related Genes Analyzed in This Study
Comprehensive list of pluripotency-related genes analyzed in RNA-seq.

| Symbol | Name | Alias |
|---|---|---|
| HNRNPL | heterogeneous nuclear ribonucleoprotein L | HNRPL, P/OKcl.14, hnRNP-L |
| HNRNPU | heterogeneous nuclear ribonucleoprotein U | DEE54, EIEE54, GRIP120, HNRNPU-AS1, HNRPU |
| HSP90B1 | heat shock protein 90 beta family member 1 | ECGP, GP96, GRP94, HEL-S-125m, HEL35 |
| HSPA8 | heat shock protein family A (Hsp70) member 8 | HEL-33, HEL-S-72p, HSC54, HSC70, HSC71 |
| HSPA9 | heat shock protein family A (Hsp70) member 9 | CRP40, CSA, EVPLS, GRP-75, GRP75 |
| HSPD1 | heat shock protein family D (Hsp60) member 1 | CPN60, GROEL, HLD4, HSP-60, HSP60 |
| IGF2BP1 | insulin like growth factor 2 mRNA binding protein 1 | CRD-BP, CRDBP, IMP-1, IMP1, VICKZ1 |
| ILF3 | interleukin enhancer binding factor 3 | CBTF, DRBF, DRBP76, MMP4, MPHOSPH4 |
| KHDC3L | KH domain containing 3 like, subcortical maternal complex member | C6orf221, ECAT1, HYDM2 |
| KLF4 | KLF transcription factor 4 | EZF, GKLF |
| LARP7 | La ribonucleoprotein 7, transcriptional regulator | ALAZS, HDCMA18P, PIP7S, hLARP7 |
| LIN28 | lin-28 homolog | CSDD1, LIN28A, LIN-28, ZCCHC1, lin-28A |
| MDN1 | midasin AAA ATPase 1 | Rea1 |
| MFGE8 | milk fat globule EGF and factor V/VIII domain containing | BA46, EDIL1, HMFG, HsT19888, MFG-E8 |
| MSH2 | mutS homolog 2 | COCA1, FCC1, HNPCC, HNPCC1, LCFS2, LYNCH1, MMRCS2, MSH-2, hMSH2 |
| MYC | MYC proto-oncogene, bHLH transcription factor | MRTL, MYCC, c-Myc, bHLHe39 |
| NANOG | Nanog homeobox | |
| NAP1L1 | nucleosome assembly protein 1 like 1 | NAP1, NAP1L, NRP |
| NCL | DnaJ heat shock protein family (Hsp40) member C5 | CLN4, CLN4B, CSP, DNAJC5A, NCL |
| NOLC1 | nucleolar and coiled-body phosphoprotein 1 | NOPP130, NOPP140, NS5ATP13, P130, Srp40 |
| NONO | non-POU domain containing octamer binding | MRXS34, NMT55, NRB54, P54, P54NRB |
| OCT4 | Octamer-binding transcription factor 4 | OCT3, POU5F1, OTF3, OTF4, OTF-3, Oct-3, Oct-4, Oct3/4 |
| PARP1 | poly(ADP-ribose) polymerase 1 | ADPRT, ADPRT 1, ADPRT1, ARTD1, PARP |

**Table S3**. List of Pluripotency-related Genes Analyzed in This Study
Comprehensive list of pluripotency-related genes analyzed in RNA-seq.

| Symbol | Name | Alias |
|---|---|---|
| PHC1 | polyhomeotic homolog 1 | EDR1, HPH1, MCPH11, RAE28 |
| PKM | pyruvate kinase M1/2 | CTHBP, HEL-S-30, OIP3, PK3, PKM2 |
| PODXL | podocalyxin like | Gp200, PC, PCLP, PCLP-1, PDX |
| POU5F1P3 | POU class 5 homeobox 1 pseudogene 3 | OCT3; OCT4; OTF3; OTF-3; OTF3L; Oct-3; Oct-4; POU5F1; POU5F1L; OCT4-pg3; POU5FLC12 |
| PRKDC | protein kinase, DNA-activated, catalytic subunit | DNA-PKC, DNA-PKcs, DNAPK, DNAPKc, DNPK1 |
| PSME3 | proteasome activator subunit 3 | HEL-S-283, Ki, PA28-gamma, PA28G, PA28gamma, REG-GAMMA |
| PTMA | prothymosin alpha | TMSA |
| REX1 | reduced expression 1 | ZFP42, REX-1, ZNF754, zfp-42 |
| RPL10A | ribosomal protein L10a | CSA19, Csa-19, L10A, NEDD6 |
| RPLP1 | ribosomal protein lateral stalk subunit P1 | LP1, P1, RPP1 |
| RPS2 | ribosomal protein S2 | LLREP3, S2 |
| RPS5 | ribosomal protein S5 | S5 |
| RPS8 | ribosomal protein S8 | S8 |
| SALL4 | spalt like transcription factor 4 | DRRS, HSAL4, IVIC, ZNF797 |
| SCD | stearoyl-CoA desaturase | FADS5, MSTP008, SCD1, SCDOS, hSCD1 |
| SEPHS1 | selenophosphate synthetase 1 | SELD, SPS, SPS1 |
| SOX15 | SRY-box transcription factor 15 | SOX20, SOX26, SOX27 |
| SOX2 | SRY-box transcription factor 2 | ANOP3, MCOPS3 |
| SRRM2 | serine/arginine repetitive matrix 2 | 300-KD, CWF21, Cwc21, HSPC075, SRL300 |
| STAT3 | signal transducer and activator of transcription 3 | ADMIO, ADMIO1, APRF, HIES |
| TCL1A | TCL1 family AKT coactivator A | TCL1 |
| TDGF1 | Teratocarcinoma-derived growth factor 1 | CRIPTO, CR, CR-1, CRGF |
| TERF1 | telomeric repeat binding factor 1 | PIN2, TRBF1, TRF, TRF1, hTRF1-AS, t-TRF1 |
| TPT1 | tumor protein, translationally-controlled 1 | HRF, TCTP, p02, p23 |
| TRIM28 | tripartite motif containing 28 | KAP1, PPP1R157, RNF96, TF1B, TIF1B |
| TUBB2B | tubulin beta 2B class IIb | CDCBM7, PMGYSA, bA506K6.1 |
| USP44 | ubiquitin specific peptidase | |
| UTF1 | undifferentiated embryonic cell transcription factor 1 | |
| YBX1 | Y-box binding protein 1 | BP-8, CBF-A, CSDA2, CSDB, DBPB |

**Table S4.** List of Endothelial Cell (EC)–related Genes Analyzed in This Study
Comprehensive list of EC-related genes analyzed in RNA-seq.

| Symbol | Name | Alias |
|---|---|---|
| ACVRL1 | activin A receptor like type 1 | ACVRLK1, ALK-1, ALK1, HHT, HHT2 |
| ADAM17 | ADAM metallopeptidase domain 17 | ADAM18, CD156B, CSVP, NISBD, NISBD1 |
| AGGF1 | angiogenic factor with G-patch and FHA domains 1 | GPATC7, GPATCH7, HSU84971, HUS84971, VG5Q |
| ANGPT1 | angiopoietin 1 | AGP1, AGPT, AGPT-1, ANG1, HAE5 |
| ANTXR1 | ANTXR cell adhesion molecule 1 | ATR, GAPO, TEM8 |
| ANXA5 | annexin A5 | ANX5, CPB-I, ENX2, HEL-S-7, PP4 |
| BSG | basigin (Ok blood group) | 5F7, CD147, EMMPRIN, EMPRIN, HAb18G |
| CCL2 | C-C motif chemokine ligand 2 | GDCF-2, HC11, HSMCR30, MCAF, MCP-1 |
| CD151 | CD151 molecule (Raph blood group) | EBS7, GP27, MER2, PETA-3, RAPH |
| CD248 | CD248 molecule | CD164L1, TEM1 |
| CD34 | CD34 molecule | |
| CDH5 | cadherin 5 | 7B4, CD144 |
| COL18A1 | collagen type XVIII alpha 1 chain | GLCC, KNO, KNO1, KS |
| COLEC12 | collectin subfamily member 12 | CLP1, NSR2, SCARA4, SRCL |
| CRADD | CASP2 and RIPK1 domain containing adaptor with death domain | MRT34, RAIDD |
| CX3CL1 | C-X3-C motif chemokine ligand 1 | ABCD-3, C3Xkine, CXC3, CXC3C, NTN |
| DCBLD2 | discoidin, CUB and LCCL domain containing 2 | CLCP1, ESDN |
| DEPP1 | DEPP autophagy regulator 1 | C10orf10, DEPP, FIG, Fseg |
| DLL4 | delta like canonical Notch ligand 4 | AOS6, delta4, hdelta2 |
| EDN1 | endothelin 1 | ARCND3, ET1, HDLCQ7, PPET1, QME |
| EDNRA | endothelin receptor type A | ET-A, ETA, ETA-R, ETAR, ETRA |
| EFNB2 | ephrin B2 | EPLG5, HTKL, Htk-L, LERK5, ephrin-B2 |
| EGFL7 | EGF like domain multiple 7 | NEU1, VE-STATIN, ZNEU1 |
| ENG | endoglin | END, HHT1, ORW1 |
| EPAS1 | endothelial PAS domain protein 1 | ECYT4, HIF2A, HLF, MOP2, PASD2 |
| EPHB4 | EPH receptor B4 | CMAVM2, HFASD, HTK, LMPHM7, MYK1 |
| EPOR | erythropoietin receptor | EPO-R |
| ESAM | endothelial cell adhesion molecule | W117m |
| FABP5 | fatty acid binding protein 5 | E-FABP, EFABP, KFABP, PA-FABP, PAFABP |
| FGF1 | fibroblast growth factor 1 | AFGF, ECGF, ECGF-beta, ECGFA, ECGFB |
| FLT1 | fms related receptor tyrosine kinase 1 | FLT, FLT-1, VEGFR-1, VEGFR1 |
| FLT4 | fms related receptor tyrosine kinase 4 | CHTD7, FLT-4, FLT41, LMPH1A, LMPHM1 |
| FN1 | fibronectin 1 | CIG, ED-B, FINC, FN, FNZ |
| ICAM1 | intercellular adhesion molecule 1 | BB2, CD54, P3.58 |
| IL13RA1 | interleukin 13 receptor subunit alpha 1 | CD213A1, CT19, IL-13Ra, NR4 |

**Table S4.** List of Endothelial Cell (EC)–related Genes Analyzed in This Study
Comprehensive list of EC-related genes analyzed in RNA-seq.

| Symbol | Name | Alias |
|---|---|---|
| IL1R1 | interleukin 1 receptor type 1 | CD121A, D2S1473, IL-1R-alpha, IL-1RT1, IL1R |
| ITGA4 | integrin subunit alpha 4 | CD49D, IA4 |
| ITGA5 | integrin subunit alpha 5 | CD49e, FNRA, VLA-5, VLA5A |
| ITGAV | integrin subunit alpha V | CD51, MSK8, VNRA, VTNR |
| ITGB1 | integrin subunit beta 1 | CD29, FNRB, GPIIA, MDF2, MSK12 |
| ITGB3 | integrin subunit beta 3 | BDPLT16, BDPLT2, BDPLT24, CD61, GP3A |
| JAG1 | jagged canonical Notch ligand 1 | AGS, AGS1, AHD, AWS, CD339 |
| JAG2 | jagged canonical Notch ligand 2 | HJ2, LGMDR27, SER2 |
| KDR | kinase insert domain receptor | CD309, FLK1, VEGFR, VEGFR2 |
| KLF4 | KLF transcription factor 4 | EZF, GKLF |
| MCAM | melanoma cell adhesion molecule | CD146, HEMCAM, METCAM, MUC18, MelCAM |
| MMP1 | matrix metallopeptidase 1 | CLG, CLGN |
| MMP2 | matrix metallopeptidase 2 | CLG4, CLG4A, MMP-2, MMP-II, MONA |
| MMP9 | matrix metallopeptidase 9 | CLG4B, GELB, MANDP2, MMP-9 |
| NECTIN2 | nectin cell adhesion molecule 2 | CD112, HVEB, PRR2, PVRL2, PVRR2 |
| NOS3 | nitric oxide synthase 3 | eNOS, ECNOS |
| NOTCH1 | notch receptor 1 | AOS5, AOVD1, TAN1, hN1 |
| NOTCH4 | notch receptor 4 | INT3 |
| NPPB | natriuretic peptide B | BNP, Iso-ANP |
| NR2F2 | nuclear receptor subfamily 2 group F member 2 | ARP-1, ARP1, CHTD4, COUPTF2, COUPTFB |
| NRP1 | neuropilin 1 | BDCA4, CD304, NP1, NRP, VEGF165R |
| NRP2 | neuropilin 2 | NP2, NPN2, PRO2714, VEGF165R2 |
| OCLN | occludin | BLCPMG, PPP1R115, PTORCH1 |
| PDGFRA | platelet derived growth factor receptor alpha | CD140A, PDGFR-2, PDGFR2 |
| PDPN | podoplanin | AGGRUS, D2-40, GP36, GP40, Gp38 |
| PECAM1 | platelet and endothelial cell adhesion molecule 1 | CD31, CD31/EndoCAM, GPIIA', PECA1, PECAM-1 |
| PGF | placental growth factor | D12S1900, PGFL, PIGF, PLGF, PlGF-2 |
| PLA2G4C | phospholipase A2 group IVC | CPLA2-gamma |
| PLAT | plasminogen activator, tissue type | T-PA, TPA |
| PLAU | plasminogen activator, urokinase | ATF, BDPLT5, QPD, UPA, URK |

**Table S4.** List of Endothelial Cell (EC)–related Genes Analyzed in This Study
Comprehensive list of EC-related genes analyzed in RNA-seq.

| Symbol | Name | Alias |
|---|---|---|
| PODXL | podocalyxin like | Gp200, PC, PCLP, PCLP-1, PDX |
| PROCR | protein C receptor | CCCA, CCD41, EPCR |
| PTGIS | prostaglandin I2 synthase | CYP8, CYP8A1, PGIS, PTGI |
| RHOB | ras homolog family member B | ARH6, ARHB, MST081, MSTP081, RHOH6 |
| RIPK1 | receptor interacting serine/threonine kinase 1 | AIEFL, IMD57, RIP, RIP-1, RIP1 |
| S1PR1 | sphingosine-1-phosphate receptor 1 | CD363, CHEDG1, D1S3362, ECGF1, EDG-1 |
| S1PR2 | sphingosine-1-phosphate receptor 2 | AGR16, DFNB68, EDG-5, EDG5, Gpcr13 |
| S1PR3 | sphingosine-1-phosphate receptor 3 | C9orf108, C9orf47, EDG-3, EDG3, LPB3 |
| SERPINE1 | serpin family E member 1 | PAI, PAI-1, PAI1, PLANH1 |
| SOD1 | superoxide dismutase 1 | ALS, ALS1, HEL-S-44, IPOA, SOD |
| SPHK1 | sphingosine kinase 1 | SPHK |
| TEK | TEK receptor tyrosine kinase | CD202B, GLC3E, TIE-2, TIE2, VMCM |
| TFPI | tissue factor pathway inhibitor | EPI, LACI, TFI, TFPI1 |
| THBD | thrombomodulin | AHUS6, BDCA-3, BDCA3, CD141, THPH12 |
| THBS1 | thrombospondin 1 | THBS, THBS-1, TSP, TSP-1, TSP1 |
| THSD7A | thrombospondin type 1 domain containing 7A | |
| TIMP1 | TIMP metallopeptidase inhibitor 1 | CLGI, EPA, EPO, HCI, TIMP |
| TNFAIP3 | TNF alpha induced protein 3 | A20, AIFBL1, AISBL, OTUD7C, TNFA1P2 |
| TNFRSF10A | TNF receptor superfamily member 10a | APO2, CD261, DR4, TRAILR-1, TRAILR1 |
| TNFRSF10C | TNF receptor superfamily member 10c | CD263, DCR1, DCR1-TNFR, LIT, TRAIL-R3 |
| TNFRSF1A | TNF receptor superfamily member 1A | CD120a, FPF, TBP1, TNF-R, TNF-R-I |
| TYMP | synthesis of cytochrome C oxidase 2 | CEMCOX1, ECGF1, Gliostatin, MC4DN2, MYP6 |
| VCAM1 | vascular cell adhesion molecule 1 | CD106, INCAM-100 |
| VEGFA | vascular endothelial growth factor A | MVCD1, VEGF, VPF |
| VWF | von Willebrand factor | F8VWF, VWD |

**Legends for Supplementary Videos**

**Video S1.** Three-dimensional confocal reconstruction showing neovascularization following transplantation of non-PAD hiPSC-ECs.

**Video S2**. Three-dimensional confocal reconstruction showing neovascularization following transplantation of PAD hiPSC-ECs.
